# A neural substrate for Bayesian integration in human parietal cortex

**DOI:** 10.1101/2021.10.30.466508

**Authors:** Nicholas M. Singletary, Jacqueline Gottlieb, Guillermo Horga

## Abstract

Making adaptive decisions often requires inferring unobservable states based on unreliable information. Bayesian logic prescribes that individuals form probabilistic beliefs about a state by integrating the likelihood of new evidence with their prior beliefs, but human neuroimaging studies on probability representations have not typically examined this integration process. We developed an inference fMRI task in which participants estimated the posterior probability of a hidden state while we parametrically modulated the prior probability of the state, the likelihood of the supporting evidence, and a monetary penalty for estimation inaccuracy. Consistent with a neural substrate for Bayesian integration, activation in left posterior parietal cortex tracked the estimated posterior probability of the solicited state and its components of prior probability and likelihood, all independently of expected value. This activation further reflected deviations in individual reports from objective probabilities. Thus, this region may provide a neural substrate for humans’ ability to approximate Bayesian inference.

## Introduction

Making adaptive decisions often requires us to infer unobservable, “hidden,” states based on probabilistic information. For example, when making a diagnosis, a physician infers an underlying illness based on observable symptoms that provide imperfect evidence for the illness. Probabilistic inference supports a variety of adaptive behaviors in humans and other animals, and deficits in probabilistic inference have been linked to psychopathology^1–4^, underscoring the importance of understanding its neural mechanisms.

According to Bayesian logic, optimal probabilistic inference requires individuals to estimate the posterior probability of a hypothesis by integrating two quantities: the *prior probability* of the hypothesis about an underlying state and the *likelihood* of new information conditional on this hypothesis, as informed by Bayes’ theorem. Although abundant evidence shows that people estimate posterior probability approximately consistently with Bayesian principles^5–8^, major questions remain about the neural mechanisms supporting the integration of prior and likelihood and their distinction from other decision variables.

A key question at the center of Bayesian inference concerns the mechanisms supporting the integration of prior and likelihood. Existing studies in humans have typically manipulated only the prior probability of a state^9^ or only the likelihood of the supporting evidence^10–14^ without manipulating the two quantities parametrically. Thus, it remains unclear how the brain combines prior and likelihood information, and to what extent neural signals reported to correlate with posterior probability instead reflect the experimentally manipulated quantity (i.e., only the prior or likelihood).

A second open question concerns the relation between posterior probability and expected value (EV). In instrumental tasks, in which participants are incentivized to accurately report the probability of a hidden state, a higher posterior probability strongly correlates with higher EV (i.e., the product of reward probability and reward magnitude). Abundant evidence shows that people can estimate probability independently of value^12,15^ and seek noninstrumental information and update their beliefs independently of rewards^16,17^. And yet, the confound between posterior probability and instrumental rewards has been largely overlooked in animal and human investigations of posterior probability^18^ or of likelihood^10,11,19^, leaving it unclear whether neural signals that have been interpreted as encoding probability are better explained as EV or other reward-related quantities.

In this study, we address these two questions using fMRI in conjunction with a novel behavioral task that required human participants to estimate the posterior probability of a hidden state based on two orthogonally manipulated variables: the prior probability of the state and the likelihood of the evidence conditional on the state. We took advantage of the brain’s separate visual modules for faces^20^ and places^21^ by yoking these states to face and place stimuli to test if any resulting BOLD activation emerged from the integration of sensory evidence from competing streams^11,22,23^. We also orthogonally manipulated the magnitude of a penalty for an incorrect estimate, incentivizing participants to provide accurate estimates while dissociating posterior probability from EV. We found that blood oxygen level– dependent (BOLD) activity in left posterior parietal cortex (PPC) not only correlated with estimates of posterior probability but also correlated separately with prior probability and likelihood independently of EV. Based on these findings, we propose this region provides a neural substrate for cognitive computations that support humans’ ability to approximate Bayesian inference.

## Results

### Probability Estimation Task

Twenty-three participants estimated the posterior probability of being in one of two states depicted as museum galleries: a *portrait gallery* that contained more pictures of faces than places or a *landscape gallery* that contained more pictures of places than faces. On each of 120 trials, participants viewed the prior probability of being in a state, the likelihood of new evidence for the state, and the potential penalty for estimation inaccuracy (**Figure 1A**). The prior probability was displayed as a percentage (e.g., 90% and 10% chance of being in the portrait and landscape galleries, respectively; **Figure 1A**). The new evidence consisted of a single sample picture that was randomly drawn from the hidden gallery and a majority-to-minority ratio indicating the likelihood (strength) of the evidence (**Figure 1A**). For example, a low majority-minority ratio (e.g., 60:40) indicated that the hidden gallery contained a relatively even mixture of images, and thus the sample picture provided weaker evidence of the hidden gallery’s identity. Conversely, a high majority-minority ratio (e.g., 90:10) indicated that the sample provided strong evidence for the hidden gallery. Then, a slider appeared along with a prompt instructing participants to report the posterior probability of being in one of the galleries (the “questioned gallery”; **Figure 1A**). The initial slider position was randomized by trial to reduce the correlation between the reported posterior probability and slider movement (**Figure S1**). Trials were divided evenly into four runs, with the questioned gallery alternating by run.

**Figure 1.**
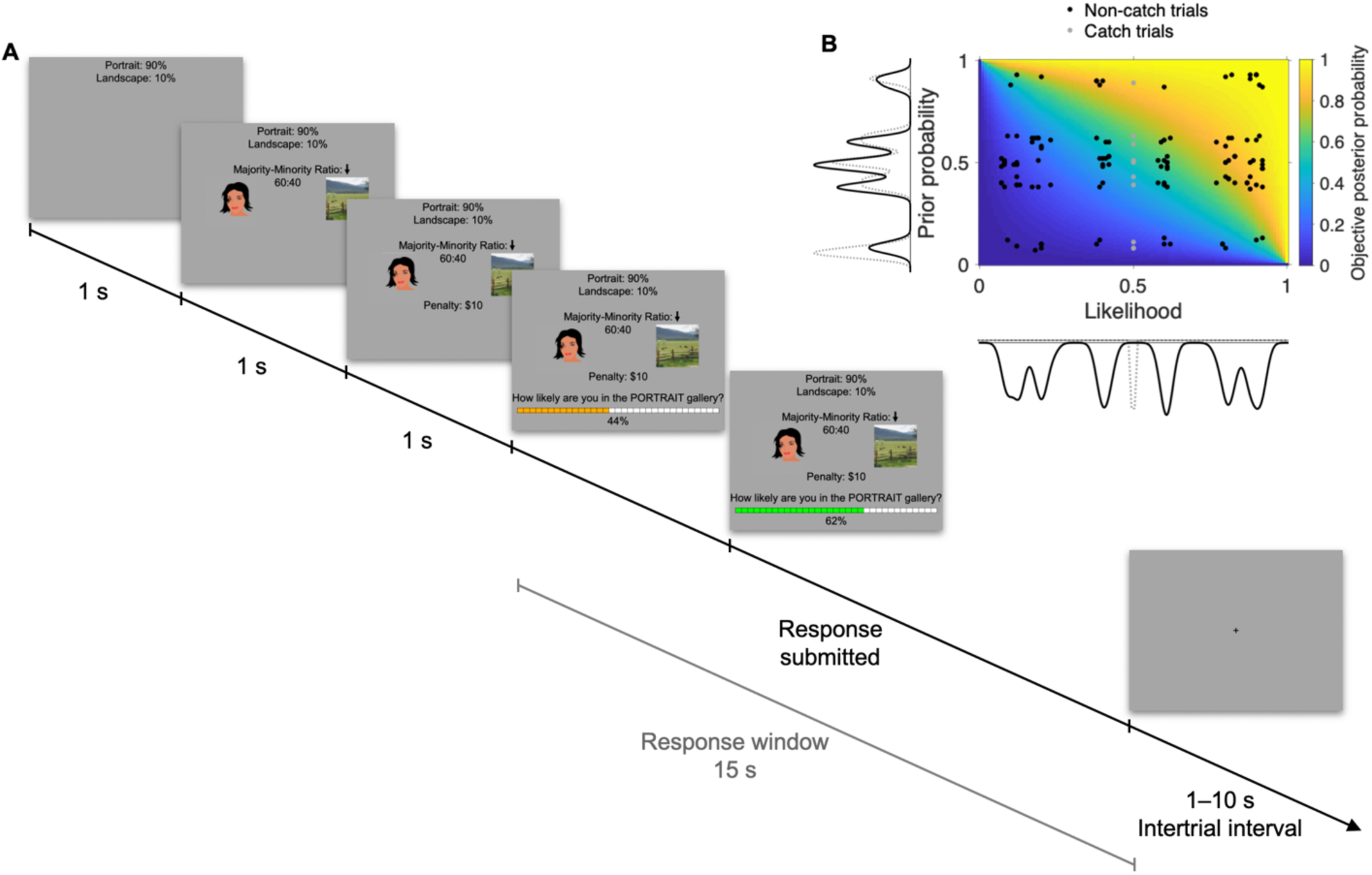
**A: Trial structure**. On each trial, participants see the **prior probability** of being in a landscape or portrait gallery (left panel, top), **one sample picture** drawn from the gallery (indicated by the arrow above it); and the **evidence strength**, represented by the ratio of majority to minority pictures in the gallery. A **decoy** picture from the opposite category is shown to control for visual activations. Together, the sample and its strength determine the **likelihood**. The **penalty** reveals how much participants could lose from their endowment due to inaccuracy in their estimate. Here, the prior probability is shown; followed by the evidence strength, sample picture, and decoy; and finally, the penalty. However, these three groups of trial elements could appear on screen in any spatial or temporal order. Face drawing for visualization; actual face stimuli were photographs of human faces (see Methods). **B:** The objective posterior probability of the questioned gallery conditional on the sample picture (colored grid) as a function of the prior probability of the questioned gallery (*y*-axis) and the likelihood of the sample conditional on the questioned gallery (*x*-axis), with points at prior-likelihood combinations that were used on trials during the scan session. To ensure that participants paid attention to the prior probability, interspersed within a session were 10 additional “catch” trials that omitted the sample picture and majority-minority ratio (see Methods); these trials were not included in the behavioral or fMRI analyses. Catch trials are in gray, and non-catch trials are in black. Because information pertaining to likelihood was missing from catch trials, the objective posterior probability is equal to the prior probability on those trials, which is equivalent to a case in which the likelihood equals 0.5. Hence, the likelihoods of the catch trials are plotted at 0.5 to reflect the omission of likelihood information on the trial.

Participants received a $30 endowment from which the inaccuracy penalty ($10 or $20) could be deducted. To prevent serial trial effects, participants were truthfully told that every trial was independent, and they did not receive feedback about their response to any trial except one trial randomly selected at the end of the session to determine their payment. The penalty on the payment trial was subtracted from the endowment with a probability equal to the squared error of the posterior estimate, providing an incentive-compatible procedure to motivate accurate probabilistic reports while dissociating posterior probability from EV (**Figure S2**; **Equation 1**, see Methods).

### Behavior

Participants experienced trials sampled from a distribution of prior probabilities and likelihoods that tiled the variable space (**Figure 1B**; **Table S1**), allowing us to distinguish the neural representations of these quantities. The reported probability estimates—henceforth, subjective posterior probabilities—increased monotonically with the objective posterior probability; when expressed as log odds (logits), there was a strong linear correlation between them (**Figure 2A**, inset; *R*^2^ = 0.77; *p* = 7.6×10^−63^), confirming that participants approximated Bayesian inference. Consistent with theory and empirical findings that decision speed increases with stronger evidence^24,25^, participants responded faster on trials where they reported higher posterior certainty (as measured by the sum of 0.5 and the absolute difference between the subjective posterior and 0.5) (**Figure 2B**; *R*^2^ = 0.46; *p* = 8.2×10^−11^).

**Figure 2.**
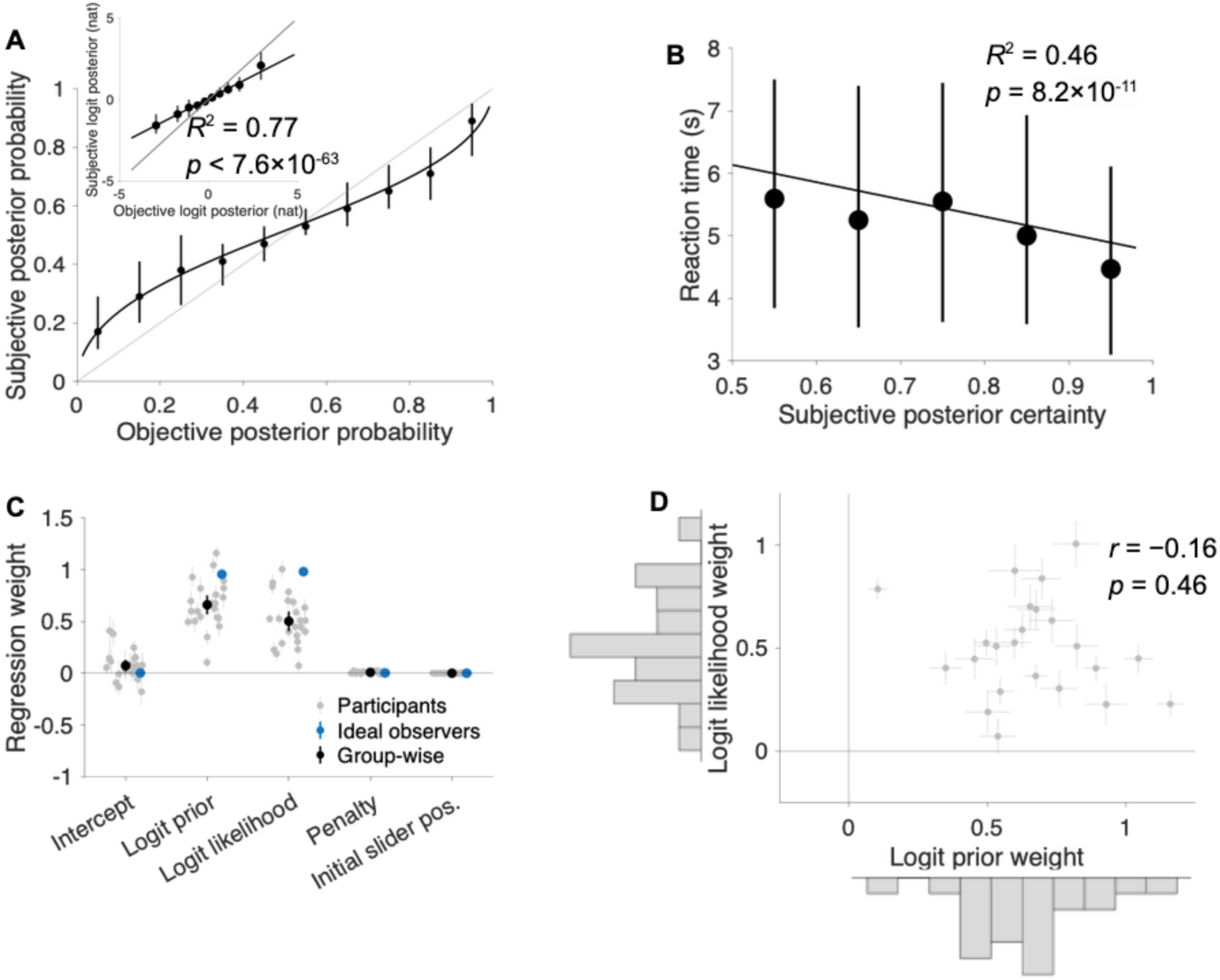
People can closely estimate posterior probability by integrating described prior probabilities and likelihoods. **A:** Plots showing a strong relationship between subjective posterior probability (reported estimates) and the objective posterior probability of the questioned gallery across all participants (*N* = 23) across non-catch trials completed during the scan session (*R*^2^ = 0.77, *p* < 0.001, linear mixed-effects model, **Equation 7**). However, subjective posterior is conservative (biased toward 0.5) compared to the objective posterior. For visualization, the median subjective posterior probability is binned by the objective posterior probability. In the inset, median subjective logit posterior is binned by objective logit posterior. The black diagonal line in the inset is the least-squares regression line. The black curve in the main panel is the regression line from the inset transformed into probability space. Error bars represent interquartile range. Gray lines represent unity. **B:** Reaction time during the scan session decreases with increasing subjective posterior certainty (*R*^2^ = 0.46, *p* = 8.2×10^−11^). Subjective posterior certainty is defined as the sum of 0.5 and the absolute difference between the subjective posterior and 0.5, such that the subjective posterior certainty is between 0.5 and 1, inclusive, with higher values indicating higher subjective certainty. For visualization, the median reaction time is binned by the subjective posterior certainty. Error bars represent interquartile range. The diagonal line is the least squares line for the individual trials. **C:** Participants incorporate prior and likelihood into their subjective posteriors, but compared to simulated ideal observers (*N* = 23), they underweight both. Regression weights during the scan session for individual participants (gray), at the group level over all participants (black), and at the group level over all simulated ideal observers (blue). Group-level coefficients estimated from a linear mixed-effects model of subjective logit posterior with mean-centered regressors (**Equation 9**, see Methods). Error bars represent 95% confidence intervals. While all terms are statistically significant at the group level, penalty and initial slider position have negligible effects compared to logit prior and logit likelihood (**Figure S4C**; **Table S5**). **D:** Scatterplot showing no significant correlation between participants’ logit prior weights and their logit likelihood weights as calculated by **Equation 9** during the scan session (Pearson correlation: *r* = −0.16, *p* = 0.46). Each point represents the logit prior and logit likelihood weight for one participant. The distribution of logit prior weights clustered just above 0.5 and the logit likelihood weights clustered near 0.5 on marginal histograms outside the *x*- and *y*-axes respectively. Error bars represent 95% confidence intervals.

To determine if participants integrated prior probability and likelihood to derive subjective posterior probability, we modeled the subjective posterior using a parameterized logit form of Bayes’ theorem that expresses subjective logit posterior as the weighted sum of the logit prior and logit likelihood (**Equation 8**; **Equation 9**). Using mixed-effects regression, the fixed-effects weights for both logit prior and logit likelihood were significantly greater than 0, indicating that participants integrated these quantities to estimate the subjective posterior (**Fig. 2C**; logit prior weight: 0.66, *T*(2,745) = 14.17, SE = 0.05, *p* = 4.7×10^−44^; logit likelihood weight: 0.50, *T*(2,745) = 10.30, SE: 0.05, *p* = 1.9×10^−24^; **Table S5**, see also **Table S4**). Yet there were specific ways in which participants deviated from Bayes’ theorem. The fixed-effects intercept was significantly greater than 0 (**Figure 2C**; intercept: 0.07, *T*(2,745) = 2.59, SE = 0.03, *p* = 0.010; **Table S5**, see also **Table S4**), indicating that participants overestimated the posterior probability of being in the questioned gallery. Also, participants underweighted logit prior and logit likelihood relative to the Bayesian ideal observers (**Figure 2C**; difference between fixed-effects logit prior weights for participants and ideal observers: −0.30, *T*(5,488) = −6.31, SE = 0.05, *p* < 0.0001; difference between fixed-effects logit likelihood weights for ideal observers and participants: −0.45, *T*(5,488) = −9.30, SE = 0.05, *p* < 0.0001), which is consistent with previous work on inference from described probabilities^8,26,27^ showing conservativism of the subjective posteriors as compared to Bayes’ theorem (**Figure 2A**). Nevertheless, integration of prior and likelihood was also apparent at the individual level, as each participant’s logit prior and logit likelihood weights were also significantly greater than 0, and they were uncorrelated across participants, supporting the independence of these two effects (Pearson correlation: −0.16, *p* = 0.46) (**Figure 2D**). Moreover, models that contained only logit prior, logit likelihood, or objective logit posterior were inferior to those that contained both logit prior and logit likelihood (**Figure S3**). Overall, these results were robust across several methods of normalizing the data and controlling for nuisance variables of penalty and initial slider position (whose effect sizes were very small; **Figure S4C**; **Table S5**), confirming that participants’ subjective posteriors were broadly consistent with Bayesian inference and justifying the use of general linear models (GLMs) to analyze task-based BOLD signals based on Bayesian variables in logit space.

A Region in Left Parietal Cortex Encodes Subjective Posterior Probability To search for candidate neural substrates of Bayesian inference, we modeled the fMRI signal during the decision period extending from the onset of the slider until the participant’s response (**Figure 1A**). First, we ran a whole-brain general linear model (WB-GLM 1) to identify regions where BOLD signals scaled positively with the subjective logit posterior of the questioned gallery. We included nuisance covariates to account for potential confounds related to subjective posterior certainty (the absolute value of the subjective logit posterior), motor preparation for hand or eye movements (initial slider position), and reward expectation (the penalty magnitude and subjective EV; **Equation 4**). This analysis revealed one candidate cluster spanning portions of the superior parietal lobule (SPL) and intraparietal sulcus (IPS) in the left posterior parietal cortex (PPC) (**Figure 3A**; **Table S6**; cluster-level FWER-corrected *p* = 0.003, permutation test). Plotting the cluster’s signal by binned subjective posterior probability confirmed that the signal increased with subjective posterior probability (**Figure 3B**).

**Figure 3.**
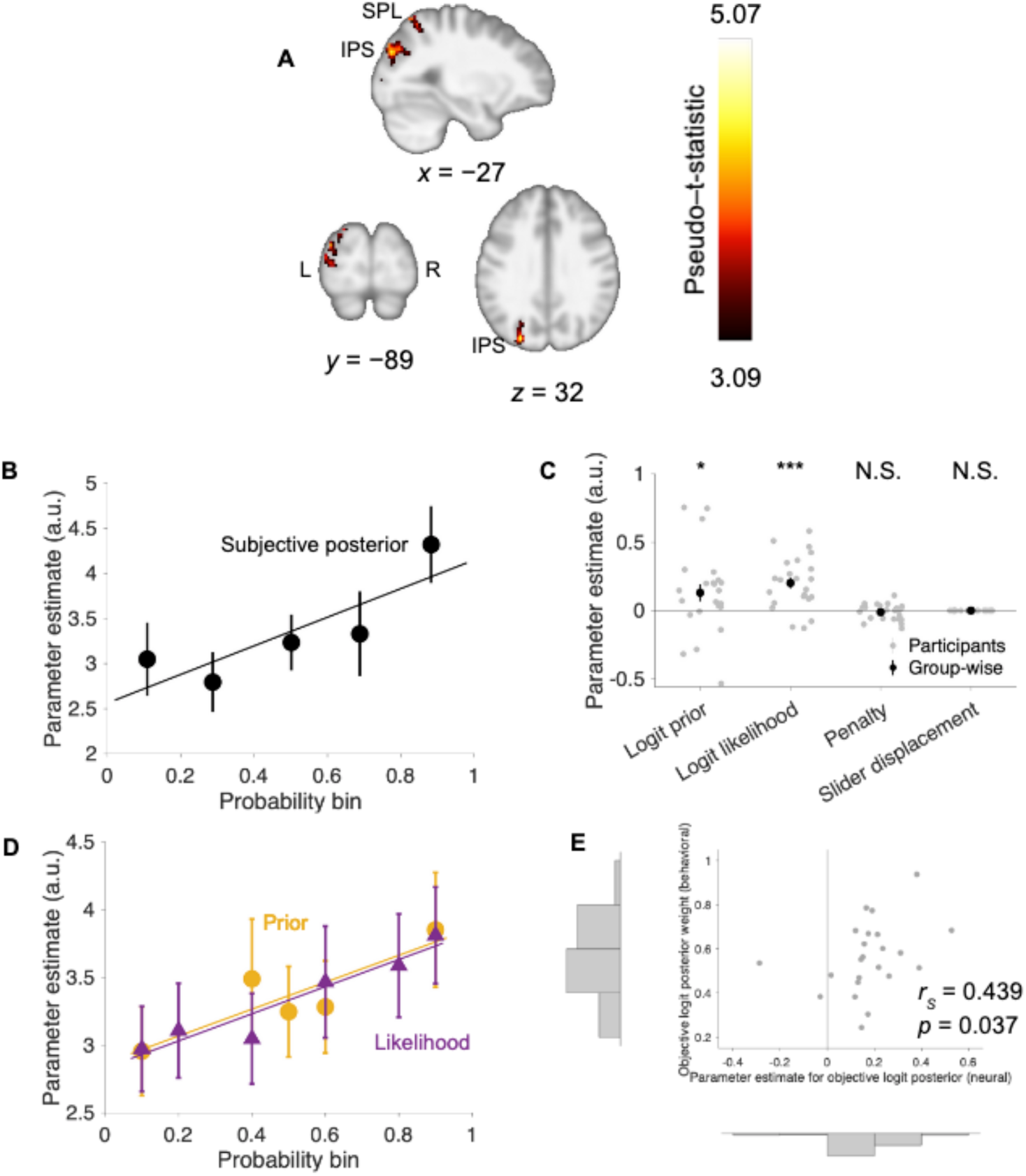
A PPC cluster encodes subjective posterior beliefs about the questioned gallery. **A:** Activation in one cluster in the left posterior parietal cortex (PPC) spanning superior parietal lobule (SPL) and intraparietal sulcus (IPS) tracks the subjective logit posterior of the questioned gallery, making it a candidate posterior belief–encoding region. Cluster-level familywise error (FWER) –corrected *p* = 0.003 (permutation test based on cluster-defining height threshold of *p* = 0.001). **B:** Corroborating results in **A**, mean activation of the cluster visually increases with binned subjective posterior probability. Points represent the mean parameter estimate for subjective posterior probability across all participants within a subjective posterior probability bin. The least-squares regression line is shown for visualization purposes for participant-wise parameter estimates as a function of posterior probability bin (individual participants’ parameter estimates not shown). From left to right, subjective posterior probability *π*(*Q*|*x*) Bin 1: 0.02 (minimum accepted subjective posterior) ≤ *π*(*Q*|*x*) < 0.2, Bin 2: 0.2 ≤ *π*(*Q*|*x*) < 0.4, Bin 3: 0.4 ≤ *π*(*Q*|*x*) < 0.6, Bin 4: 0.6 ≤ *π*(*Q*|*x*) < 0.8, Bin 5: 0.8 ≤ *π*(*Q*|*x*) ≤ 0.98 (maximum accepted subjective posterior). Error bars represent standard error. Despite the appearance of a slight nonlinearity among the data points, a quadratic term for the subjective logit posterior is not statistically significant after controlling for the linear term (**Table S7**). **C:** Post-hoc analyses of the cluster in **A** show it is significantly positively associated with both the logit prior of the questioned gallery and the logit likelihood of the sample picture conditional on the questioned gallery after accounting for the inaccuracy penalty and slider displacement on the trial. Error bars represent standard error. *: *p* < 0.05, ***: *p* < 0.001. **D:** Corroborating the significantly positive linear effects of logit prior and logit likelihood in the cluster, mean activation increases with binned prior probability (yellow) and likelihood (purple). Points represent mean parameter estimate for prior probability and likelihood across all participants within a prior probability or likelihood bin, respectively. Respective trendlines are the least squares lines for participant-wise parameter estimate as a function of prior probability and likelihood bins (individual participants’ parameter estimates not shown). From left to right, prior probability Pr(*Q*) Bin 1: 0.07 ≤ Pr(*Q*) ≤ 0.13, Bin 2: 0.37 ≤ Pr(*Q*) ≤ 0.43, Bin 3: 0.47 ≤ Pr(*Q*) ≤ 0.53, Bin 4: 0.57 ≤ Pr(*Q*) ≤ 0.63, Bin 5: 0.87 ≤ Pr(*Q*) ≤ 0.93. From left to right, likelihood Pr(*x*|*Q*) Bin 1: 0.07 ≤ Pr(*x*|*Q*) ≤ 0.13, Bin 2: 0.17 ≤ Pr(*x*|*Q*) ≤ 0.23, Bin 3: 0.37 ≤ Pr(*x*|*Q*) ≤ 0.43, Bin 4: 0.57 ≤ Pr(*x*|*Q*) ≤ 0.63, Bin 5: 0.77 ≤ Pr(*x*|*Q*) ≤ 0.83, Bin 6: 0.87 ≤ Pr(*x*|*Q*) ≤ 0.93. Error bars represent standard error. **E:** Parameter estimate for BOLD signal to objective logit posterior (*x*-axis) within the PPC cluster is positively correlated with behavioral objective logit posterior weight (*y*-axis) across all participants (Spearman correlation: 0.439, *p* = 0.037), suggesting that distortions in neural representations of posterior probability in PPC contribute to the degree of distortion in participants’ subjective posterior probabilities. Each point represents one participant.

To determine if this response represented Bayesian integration, we analyzed the average activity in this cluster with a GLM that had separate terms for logit prior and logit likelihood (alongside nuisance covariates for penalty and slider displacement; fROI-GLM 1). Activation in the PPC cluster had significant positive associations with both the logit prior (parameter estimate: 0.131, *T*(42,960) = 2.070, SE = 0.063, *p* = 0.039) and logit likelihood (parameter estimate: 0.202, *T*(42,960) = 5.124, SE = 0.039, *p* < 3.0×10^−7^), consistent with the effect of subjective posterior, but not with slider displacement (parameter estimate: 5.9×10^−5^, *T*(42,960) = 0.015, SE = 0.004, *p* = 0.988) or penalty (parameter estimate: −0.011, *T*(42,960) = −0.953, SE = 0.012, *p* = 0.341) (**Figure 3C**; **Table S8**). Visualization of the binned data (**Figure 3D**) confirmed that this cluster’s signal increased with the prior and likelihood, consistent with Bayesian integration. Analyses to further examine a unique contribution of prior probability or likelihood while controlling for subjective posterior yielded nonsignificant results (**Figure S5**; **Table S9**; **Table S10**), likely due to collinearity between these variables.

Next, we evaluated whether interindividual differences in neural probability signals could explain differences in behavior. To do so, we first analyzed the average activation within the PPC cluster using a GLM with a term for the *objective* logit posterior and nuisance regressors for penalty and slider displacement (fROI-GLM 2), finding that the neural parameter estimates and behavioral weights for objective logit posterior positively correlated across individuals (Spearman correlation: 0.439, *p* = 0.037; **Figure 3E**; **Figure S6A**). This correlation suggests that distortions in PPC activation with respect to the objective posterior probability contribute to the degree of the observed conservatism of participants’ reported estimates (**Figure 2A**). Logit prior and logit likelihood signals (as determined by fROI-GLM 1) were not significantly correlated with their respective behavioral weights (Spearman correlation for logit prior: 0.008, *p* = 0.973; Spearman correlation for logit likelihood: 0.156, *p* = 0.475; **Figure S6B–C**).

Whole-brain analyses did not reveal additional clusters exhibiting significant correlation between activation tracking objective logit posterior, logit prior, or logit likelihood and participants’ respective probability weights. Furthermore, to identify clusters showing a significant effect of the *objective* logit posterior, we defined a whole-brain GLM (WB-GLM 2) that contained the same nuisance regressors as WB-GLM 1 but replaced the terms for subjective logit posterior and its absolute value (subjective posterior certainty) with the corresponding terms for the objective posterior according to Bayes’ theorem; this yielded no clusters with a significant effect of objective logit posterior. Combined with the findings indicating activation tracking subjective posterior and neurometric-psychometric match in the distortion of posterior, this suggests that neural probability signals are indeed subjective (leniently thresholded results in **Figure S7**).

To verify that we did not miss additional candidate regions for prior-likelihood integration, we defined another whole-brain GLM to identify clusters showing a conjunction of logit prior and logit likelihood effects (WB-GLM 3). WB-GLM 3 also contained the same nuisance regressors as WB-GLM 1 but instead replaced the terms for subjective logit posterior and subjective posterior certainty with corresponding terms for logit prior, prior certainty, logit likelihood, and likelihood certainty. Despite using a lenient threshold (uncorrected cluster-forming *p*-value threshold: 0.05), this analysis did not reveal clusters with a significant conjunction of prior and likelihood effects (**Figure S8**).

### Lack of Encoding of Category-Concordant Subjective Posterior Probabilities in Face- and Place-Selective Areas

Theoretical and empirical work suggests that inferring the probabilities of underlying states engages areas selective to sensory evidence associated with those states^11,23,28^. To determine if this was the case on our task, we used an independent face-place localizer (see Methods) to identify participant-specific face- and place-selective functional regions of interest (fROIs) (**Figure 4A**) and examined if they represented subjective posterior probability of the portrait and landscape galleries or its components. To facilitate this analysis, we used fROI-GLM 1 to measure prior and likelihood activation and defined another GLM (fROI-GLM 2) to measure activation tracking subjective logit posterior. (fROI-GLM 2 was the same as fROI-GLM 1 but replaced fROI-GLM 1’s regressors for logit prior and logit likelihood with one for subjective logit posterior.) The appropriate contrasts (see Methods) revealed no significant activation tracking category-concordant subjective logit posterior (**Figure 4B; Figure S9A**; **Table S11**) or logit prior (**Figure S9B**; **Table S12**) within the face and place fROIs. While activation in the face fROI significantly tracked the logit likelihood conditional on the portrait gallery (parameter estimate: 0.110; *T*(853,978) = 3.400, SE = 0.032, *p* < 0.001), the place fROI was did not show the analogous response to the logit likelihood conditional on the landscape gallery (**Figure S9C**; parameter estimate: −0.015; *T*(853,978) = −0.384, SE = 0.038, *p* = 0.701). We also investigated the possibility that face and place fROIs encoded probability more strongly if the questioned gallery was concordant with the fROI’s preferred category. However, analyses that divided runs by the questioned gallery revealed no effect of the concordance between the questioned gallery and preferred category on either fROI’s tracking of subjective logit posterior or its components (test statistic for interaction between cluster and fMRI contrast in ANOVA for subjective logit posterior: *F*(3) = 0.81, *p* = 0.41; **Figure 4C**; **Figure S9D**; **Table S13**; **Table S14**; test statistic for interaction between cluster and fMRI contrast in ANOVA for logit prior and logit likelihood: *F*(3) = 0.76, *p* = 0.58; **Figure S9E–F**; **Table S15**; **Table S16**). Finally, whole-brain analyses of activation tracking subjective logit posterior (WB-GLM 1) and by logit prior and logit likelihood (WB-GLM 3) with respect to the portrait and landscape galleries instead of the questioned gallery produced mostly nonsignificant results (**Table S17**; **Table S18**; **Figure S10–Figure S11**). Together, these findings suggest that probabilistic information in this task was encoded relative to the questioned gallery and likely not in a category-specific format.

**Figure 4.**
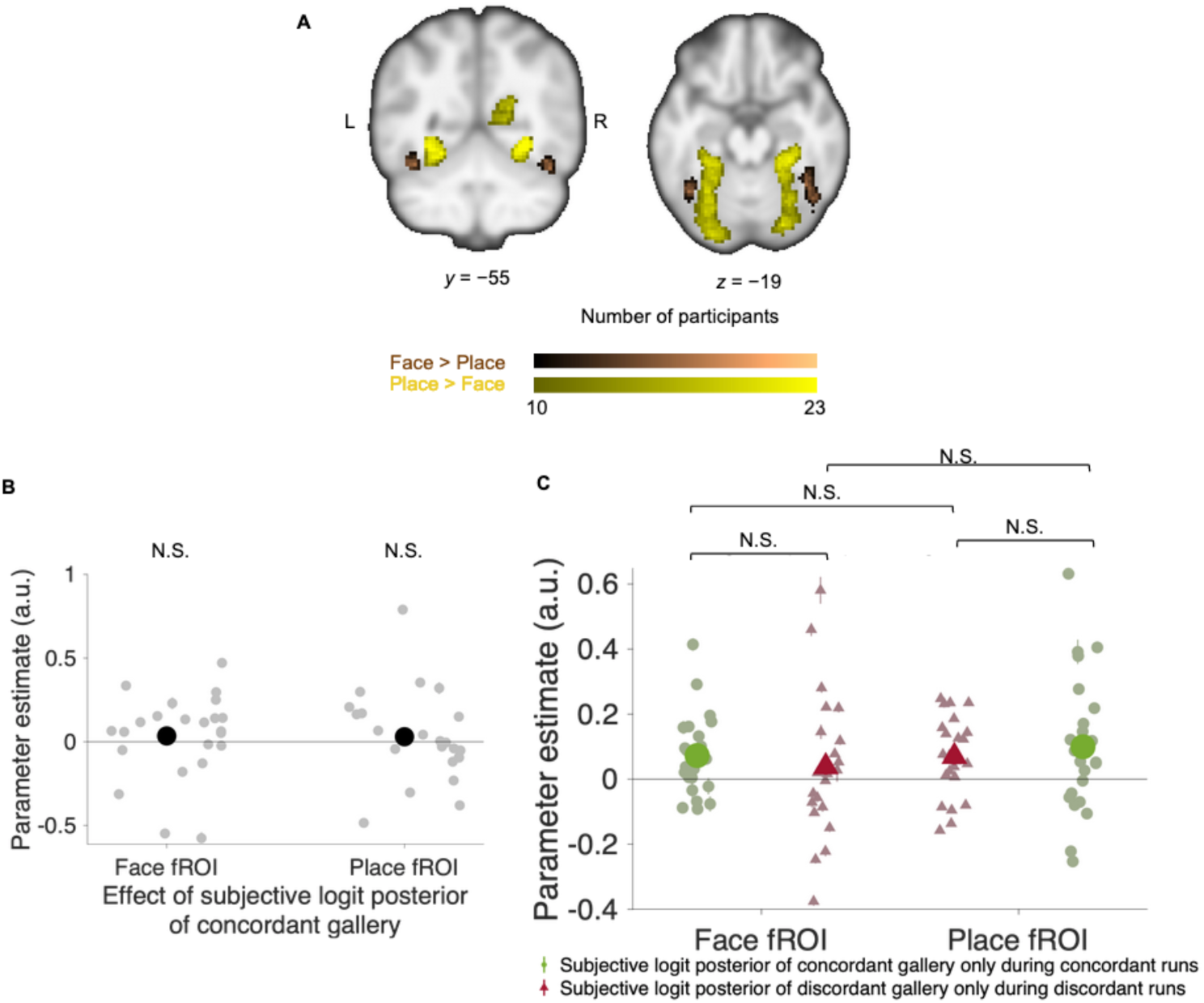
Lack of evidence that face- and place-selective functional regions of interest (fROIs) encode subjective posterior beliefs about category-concordant galleries. **A:** Participants’ face- (brown) and place-selective (yellow) functional regions of interest (fROIs), normalized to MNI space for visualization purposes. Within subjects, voxels defining fROIs were selected in native space based on an uncorrected *p*-value threshold of 0.001 for the respective contrasts (Face>Place or Place>Face) from an independent functional localizer task in a search constrained to the occipital and temporal lobes. Face-selective fROIs encompass the fusiform face area and place-selective fROIs encompass the parahippocampal place area. **B:** Neither the face-selective nor the place-selective regions show significant effects of the subjective logit posterior of their concordant galleries (i.e., the galleries corresponding to the “preferred” stimuli of that region: portrait gallery corresponding to face fROI and landscape gallery corresponding to place fROI). Because there were only two options (portrait gallery or landscape gallery), the posterior probabilities of the two galleries are complementary. Group-level statistics in black while participant-level statistics in gray. **C:** After dividing trials by their questioned galleries (portrait or landscape), neither fROI showed preferential activation tracking the posterior probability of its concordant gallery and neither posterior probability had a higher parameter estimate in its concordant fROI. Group-level statistics in saturated colors while participant-level statistics in pastel colors. Error bars represent standard error. On most points, the error bars are too small to be visible. N.S.: not significant.

## Discussion

To elucidate neural substrates for Bayesian inference, we designed a new fMRI task in which participants were incentivized to report accurate estimates of posterior probabilities of two alternative hidden states based on information about their prior probabilities, likelihoods, and an inaccuracy penalty. fMRI analyses revealed a cluster encompassing the SPL and IPS in left PPC that tracks subjective posterior probability and its components of prior probability and likelihood in ways that could not be attributed to other signals commonly localized to PPC, such as visuospatial cognition^29,30^ or expected value^31^. Neural and behavioral sensitivity to the objective posterior probability were correlated across individuals, suggesting that PPC modulates individual susceptibility to distortions in probabilistic inference. These results add to our understanding of the neural mechanisms of probabilistic inference and highlight the PPC as a candidate substrate for the integration of prior and likelihood into a subjective representation of posterior probability.

Our study goes beyond most existing approaches to identifying neural probabilistic representations by explicitly localizing a representation of the posterior probability and confirming that this representation incorporated both algebraic components of posterior probability—prior probability and likelihood. This approach is inspired by “axiomatic” approaches to identifying the representations of distinct quantities that comprise reward prediction error^32,33^. As such, our findings go beyond previous neuroimaging studies that manipulated either the prior probability or the likelihood in isolation, reporting activations tracking the manipulated quantity in PPC^12,14^ or in other regions^9,11–13^. The relatively coarse spatial resolution of fMRI does not allow us to conclude whether prior and likelihood information were integrated in the PPC cluster instead of being independently encoded by adjacent neuronal populations. However, the section of the PPC cluster in IPS overlaps with the human homologue to monkey lateral intraparietal (LIP) area^34–36^, which has been associated with probabilistic inference in support of decision-making^19,37^ and was recently shown contain separate representations of likelihood^38^ and prior uncertainty^39^. Thus, our findings reinforce a key role of the PPC in Bayesian inference and motivate continued studies of Bayesian integration in individual neurons.

Our study also goes beyond existing approaches in controlling for a confound between probability and expected value (EV). In instrumental tasks where participants are incentivized to infer the identity of a hidden state^11^, a higher posterior probability of being in a state correlates with a higher probability of being correct—and rewarded—for reporting that state. Previous work using instrumental tasks^18^ shows that EV signals in value-encoding brain regions contain contributions from prior probability and likelihood, further emphasizing the importance of isolating upstream posterior signals. In our task we ruled out an EV confound by referring to value-neutral states (e.g., asking participants to estimate the probability of a portrait or landscape gallery instead of the probability of a more or less desirable state) and by independently manipulating the magnitude of a penalty for estimation inaccuracy. Our conclusion that the PPC cluster represents posterior probability independently of EV is consistent with a previous study in which the PPC represented the probability but not the expected value of a prize^12^. It is also consistent with recent results that monkey LIP neurons encode stimulus likelihood and prior uncertainty independently of instrumental rewards^38–40^. Importantly, while our results control for EV, they do not speak against the notion that PPC can encode reward quantities, as found in other studies in monkeys^31^ and humans^41^. Thus, our findings suggest that human PPC can represent probabilistic information independently of EV.

Previous studies implicated monkey LIP neurons in the integration of probabilistic information from noisy sensory features (e.g., a random dot motion display)^19,42^ and from abstract predictive cues whose likelihoods are learned from experience^37,43,44^. By contrast, our study conveyed information about priors and likelihood through verbal and numeric *description*. Combined with previous work, our results thus suggest that PPC is recruited in probabilistic inference across tasks that convey probabilistic information in different ways. Supporting this view, in our study, posterior probability signals in PPC were referenced to the questioned gallery (i.e., the gallery whose posterior probability participants were asked to report) and not to representations of the possible hidden states in face- or place-selective areas, as may have been expected during integration of sensory evidence from competing streams^11,22,23^. This result may be due to our task design, in which we alternated the questioned gallery in a blocked, predictable manner to reduce switching costs; it is possible that, had we presented the sample picture before specifying the question, we would have uncovered place- and face-concordant probabilistic encoding. But regardless of these potential alternative outcomes, our findings show that probabilistic beliefs may be represented with respect to the task objective and suggest a domain-general involvement of PPC in probabilistic inference.

The abstract nature of description-based inference, in turn, raises questions about the relationship between our findings and human IPS activations when people solve arithmetic problems through approximation or exact calculation^45,46^. The activations we find are unlikely to reflect the mere addition of the logit prior and likelihoods because arithmetic-related IPS activation increases with problem complexity^46^, which does not covary with posterior probability. However, an interesting topic for future research concerns the mechanisms of probabilistic computations based on numeric cues and their relation to arithmetic abilities, especially since the perception of risk and probabilities greatly differs when probabilities are conveyed through description instead of being learned from experience^27,47^.

In sum, we show that a region of PPC encodes the subjective posterior probability of a hidden state by integrating prior and likelihood in a representation separable from visuospatial variables and expected value. Together with previous literature, our results support the notion that PPC may perform domain-general probabilistic inference. We also find that PPC may contribute to individual variability in distortions in probabilistic inference, suggesting a possible role in inferential psychopathology and its treatment that warrants further study.

## Methods

### Participants

Forty-four healthy, right-handed participants (17 female) were recruited through fliers posted on the Columbia University campus and through the recruitment system for the Columbia Business School Behavioral Research Lab. This pool consisted of Columbia University students, other Columbia affiliates, and affiliates of other universities in the New York Metropolitan Area, and they did not report any psychiatric or neurological disorders. Participants first completed a session outside of the scanner (prescan session); 14 participants were not allowed to advance to the scan session because their responses during the prescan session reflected disengagement or lack of comprehension (see “Performance-Based Exclusion Criteria”). Another participant was excluded because of excessive motion inside the MRI scanner, and 6 participants withdrew from the study. As a result, the final sample consisted of 23 participants (8 female). Experimental procedures were approved by the Columbia University Institutional Review Board, and all participants provided signed informed consent.

### Experimental Tasks and Sessions

The full study took place over a prescan and a scan session scheduled on different days. Both sessions consisted of the Probability Estimation Task (the primary behavioral task in this study; **Figure 1A**) and an information-sampling task outside of the scope of this study. The scan session additionally included a Face-Place Localizer Task. We wrote all tasks in MATLAB using the Psychtoolbox extensions^48,49^.

### Prescan Session

The prescan session was administered on a computer outside of the scanner. Participants viewed a narrated slideshow on the instructions for the Probability Estimation and information-sampling tasks before starting each task. They were also administered comprehension quizzes on the instructions, which they had to pass before proceeding (see “Performance-Based Exclusion Criteria”). After passing the instructions quiz, participants completed 10 practice trials of the Probability Estimation Task to familiarize them with its incentivization structure while avoiding overtraining. Each practice trial was followed by a corresponding mock payout trial to show participants what they could have earned from that trial in the main task based on their submitted estimate if the trial had been chosen for payout; however, these practice trials did not affect the participants’ earnings. Then, participants completed the Probability Estimation Task, after which their performance was evaluated to determine if they met the remaining performance criteria to advance to the scan session; if not, they were removed from the study.

### Scan Session

Participants watched a summarized version of the instructions slideshows before completing the information-seeing task (not used in this study), the Probability Estimation Task, and the Face-Place Localizer in the MRI scanner. Participants were debriefed at the end of the session.

### Probability Estimation Task

The Probability Estimation Task consisted of an *Estimation Stage* followed by a *Payout Stage*. To encourage participants to remain engaged with the task, we designed the task so that participants’ estimation accuracy influenced their earnings. During the Estimation Stage, participants estimated the posterior probability of a hidden state depicted as a museum gallery. During the Payout Stage, one trial was drawn at random to determine the participant’s payout. At the beginning of each session, the participant was given a $30 endowment from which a penalty of $10 or $20 would be withdrawn during the Payout Stage depending on the deviation of the participant’s estimate from the eventual outcome. We based participants’ earnings on a single estimation trial instead of averaging potential earnings across all estimation trials to discourage participants from allowing their accuracy to decline during later trials if they had believed their performance on earlier trials had been sufficient to make high earnings.

### Estimation Stage

The Estimation Stage of the Probability Estimation Task consisted of 130 trials divided into 4 runs of 32, 33, 32, and 33 trials, respectively. On each trial, participants had to estimate the posterior probability of being in either a *portrait gallery* that contained more pictures of faces than places or a *landscape* gallery that contained more pictures of places than faces. Participants viewed the *prior probability* of being in each gallery and possibly also the *likelihood* of the sample picture. On 10 “catch trials” distributed randomly through the Estimation Stage, the sample picture and likelihood were absent, so participants would have to estimate the posterior probability with the prior probability only (**Figure 1B**).

#### Trial display

The prior probability was displayed as a percentage (e.g., 90%). The likelihood information consisted of one face picture^50^, one place picture^51^, and potentially the majority-to-minority ratio of pictures in the hidden gallery (e.g., 60:40). One face picture and one place picture were always shown on each trial to control for the fMRI activation by the appearance of faces and places, as we were instead interested in the degree of potential face- and place-selective activation by probabilistic information. During non-catch trials, the likelihood would consist of a majority-minority ratio of picture types in the hidden gallery, one sample picture randomly drawn from the hidden gallery, and one decoy picture which signaled the opposite category from the sample picture (i.e., if the sample picture were a face, the decoy would be a place and *vice versa*) (**Figure 1A**). An arrow appeared over the true sample picture so that participants could distinguish it from the decoy picture (**Figure 1A**). During catch trials, in place of the likelihood, there were two decoy pictures and no majority-minority ratio (not shown). Participants were also shown the *penalty* that they could lose from the endowment if the trial were chosen for payout (see “Payout Stage”).

A trial began with the prior probability, likelihood information, or penalty appearing over a gray background (**Figure 1A**). The prior probability, likelihood (or likelihood decoy), and penalty appeared one at a time with the first component appearing at the instant of trial start and the succeeding components following the previous component by 1 s (**Figure 1A**). The trial components’ spatial order of appearance was stable throughout the prescan and scan sessions but counterbalanced by participant so that participants could expect the information to be in the same place while allowing us to control for potential effects of spatial order. The trial components’ temporal order of appearance was randomized by trial to control for potential primacy and recency effects.

#### Submission

Participants completed a trial by reporting their estimate of the posterior probability of the questioned gallery (the gallery in the prompt below the slider) by using a trackball to move a slider that appeared at the bottom of the screen 1 s after the last trial component. The slider remained on screen for 15 s (“response window,” **Figure 1A**). We chose a response window of 15 s because it was the shortest response window that captured approximately 80 percent of repsonses from 80 percent of participants during piloting. The selected posterior probability estimate was indicated by the amount of the slider from left to right that was highlighted in orange and by an explicit percentage below the slider. Both these indicators were updated in real time. To account for potential framing effects induced by the prompt, the questioned gallery was the portrait gallery on the first and third runs while it was the landscape gallery on the second and fourth runs. The slider was divided into 33 discrete posterior probability bins, increasing in steps of 3 percent from 2 percent on the left to 98% on the right. We chose these increments to discourage participants from anchoring to “round” numbers (e.g., multiples of 10% or 25%) and so that submitted posterior estimates could not be 0% or 100%, which would make the behavioral model inestimable (see “Modeling Subjective Posterior”). The participant confirmed their response by clicking a button on the trackball, after which the highlighted section of the slider would change colors from orange to green to indicate that the response had been recorded. The screen remained frozen until the end of the response window plus 0.5 s. If the participant did not submit a posterior probability estimate within the 15-s response window, instead, the slider would freeze for 0.5 s and the percentage below the slider would be replaced by text reading, “Estimate not submitted” (not shown). To encourage participants to respond within the response window, participants were truthfully warned that if a response were missing from a trial that happened to be chosen for payout, they would automatically lose that trial’s penalty. Across all participants during the scan session, only 10 trials were missed (all non-catch trials), with 3 participants missing one trial, 2 participants missing two trials, and 1 participant missing three trials.

#### Intertrial interval

Each estimate trial was followed by an intertrial interval during which a small, black fixation cross appeared over the gray background (**Figure 1A**). To maximize the efficiency of parameter estimation for the general linear models in the fMRI analysis, the duration of each intertrial interval was drawn from an exponential distribution with mean 3.5 s, truncated with a lower bound of 1 s and an upper bound of 10 s^52^.

#### Preventing serial trial effects

Since the task was designed to investigate prior-likelihood integration after receiving only one sample, we sought to prevent behavioral artifacts from serial trial effects such as the gambler’s fallacy. Therefore, we truthfully told participants that each estimation trial was independent from all other estimation trials, and the identity of a trial’s hidden gallery was never revealed during the Estimation Stage.

#### Selection of parameters for estimation trials

To determine the set of prior probabilities and majority-minority ratios used for the non-catch trials in each session, we randomly sampled 60 trials from discrete bins that we established for prior probability (0.1, 0.4, 0.5, 0.6, and 0.9, arbitrarily chosen as the prior of the portrait gallery) and majority-minority ratio (60:40, 80:20, and 90:10). Majority-minority ratios represented evidence strength, which was defined on the interval (0.5,1] and corresponded to the numerator of the majority-minority ratio divided by 100.

To sample the bins of prior probability and evidence strengths for each session, we used the following method. While equiprobably sampling each prior– evidence strength combination would have maximized the statistical power to detect the influence of prior and likelihood on the posterior estimates, or subjective posteriors, the Probability Estimation Task was complemented in sessions by an information-sampling task (not shown) on which the expected information gain of a sample picture was an important variable. To maximize power to detect effects of expected information gain on the complementary task would have required biased selection of trials with high evidence strength (likelihood close to 0 or 1) and low prior certainty (prior probability close to 0.5) to yield a uniform distribution of expected information gains. To make the distribution of priors and evidence strengths commensurate between the Probability Estimation and information-sampling tasks, we sampled prior–evidence strength combinations in a way that addressed the demands of both tasks: half the trials sampled prior–evidence strength combinations equiprobably to maximize statistical power on the Probability Estimation Task, and the other half sampled prior–evidence strength combinations in a biased manner to maximize statistical power for the information-sampling task.

A random jitter (−0.03, −0.02, −0.01, 0, 0.01, 0.02, or 0.03) was then added to each prior probability and evidence strength with equal probability. A “true” hidden gallery was assigned to each trial based on the prior probability of the portrait gallery (e.g., if the prior probability was 0.6, there was a 60% chance the trial’s hidden gallery would be a portrait gallery and a 40% chance it would be a landscape gallery). A trial’s sample picture was assigned to signal the hidden gallery with a probability equal to the trial’s evidence strength (e.g., there was a 60% chance that the sample would be a face on a trial on which the hidden gallery was the portrait gallery and the evidence strength was 0.6). These 60 trials were duplicated for each condition of inaccuracy penalty ($10 or $20). The parameters for the remaining 10 catch trials were assigned by assigning two trials to each of the five prior probability bins (one trial for each penalty condition) and jittering the prior probabilities by the aforementioned jittering method. The order of the trials was then randomly permuted, and the session was separated into four runs, with 32 trials in the first and third runs and 33 trials in the second and fourth runs. **Table S1** contains a list of parameters for each estimation trial in the scan session. **Figure 1B** displays the prior-likelihood combinations for the scan session, with the result of the binning and jittering process visible as peaks on the kernel density plots against each axis.

### Payout trial

#### Trial

After the Estimation Stage was complete, one estimation trial was chosen at random with equal probability to determine the participant’s payment. This trial was displayed along with its reported posterior probability estimate from the Estimation Stage. If the participant had failed to report a posterior probability estimate on that trial, the participant was notified that the inaccuracy penalty would be automatically subtracted from their endowment, and the session would end. Otherwise, the trial’s hidden gallery was revealed, and the participant was told whether they would keep all their endowment or if they had lost the error penalty, depending on the posterior probability estimate that they had submitted during the Estimation Stage.

#### Binarized scoring rule

The probability that the participant would lose the penalty was determined by a binarized scoring rule with a quadratic loss function ^53^. The binarized scoring rule incentivizes the participant to accurately estimate the posterior probability by increasing the probability that the participant would lose the penalty with increasing error between the posterior estimate and the identity of the trial’s hidden gallery. The outcome was binary—either the participant lost the full penalty or they did not—allowing the scoring rule to account for differing valuations of the loss due to differing risk preferences among participants^53^. The probability *p*_*loss*_ that the participant would lose the penalty based on their reported subjective posterior probability of the questioned gallery *π*(*Q*| *x*)is given by **Equation 1**.

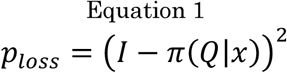

Here, *I* = 1 if the hidden gallery is the questioned gallery, and *I* = 0 if the hidden gallery is not the questioned gallery. The expected value *V* of the hidden gallery is given by **Equation 2**, where *E* is the endowment, *W* is the penalty, and Pr(*Q*|*x*)is the objective posterior probability of the questioned gallery. This relationship is plotted in **Figure S2A–B**.

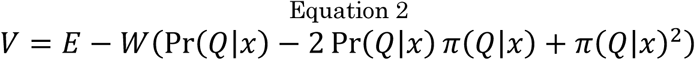

*p*_*loss*_ is minimized by the objective posterior probability (**Equation 5**), thus maximizing the expected value of a trial. For an ideal observer who submits the exact objective posterior probability, the expected value *V*_*ideal*_ of an estimation trial is given by **Equation 3**.

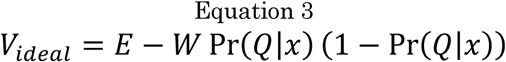

Since the Probability Estimation Task only accepts reported probabilities in bins (**Figure 1A**), on the real task, *p*_*loss*_ is minimized by reporting a subjective posterior as close as possible to the objective posterior. Assuming that participants believe that their reported subjective posteriors are equal to the objective posteriors, we can calculate the subjective expected value *V*_*subjective*_ by replacing the objective posterior probability in **Equation 3** with the subjective posterior probability (**Equation 4**).

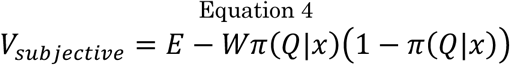

#### Face-Place Localizer

To localize face- and place-selective visual modules, we included a face-place functional localizer during the scan session. During the localizer task, participants viewed a picture of a face or a place on a gray background for 1 s, followed by a fixation cross for 1/3 s. Stimuli were blocked by type (face or place); each block consisted of 12 presentations of the same picture category followed by a rest period of 16 1/3 s. Participants completed two runs of the Face-Place Localizer. Each run consisted of 10 blocks. The Face-Place Localizer was administered as a one-back task: participants had to right-click on a trackball if the picture on screen was the same as the previous picture while they had to left-click if the picture on screen was different from the previous picture.

#### Image Sets

The same image sets were used in the Probability Estimation Task and the Face-Place Localizer. Images of faces were selected from the CNBC Faces database by Michael J. Tarr, Center for the Neural Basis of Cognition and Department of Psychology, Carnegie Mellon University, http://www.tarrlab.org, funded by NSF award 0339122 and used in Righi et al.^50^. Images of places were selected from the database for Konkle et al.^51^, available from the Computational Perception and Cognition Lab at MIT (http://olivalab.mit.edu/MM/sceneCategories.html).

#### Performance-Based Exclusion Criteria

To ensure participant comprehension and engagement during the scan session, we assessed participants’ performance during the prescan session before we allowed them to advance to the scan session. Participants had to meet the criteria for both the Probability Estimation Task and the information-sampling task to advance to the scan session, which were

1. Task comprehension: Participants had to correctly answer at least 80 percent of the question on the comprehension quizzes for the Probability Estimation and the information-sampling tasks.
2. Task completion: Participants could miss no more than 6 percent of trials on either task.
3. Minimal sensitivity to prior probability: On catch trials of the Probability Estimation Task (trials without a sample), the Pearson correlation between reported subjective posterior and the objective posterior must have been at least 0.89 (*α* = 0.05).
4. Minimal sensitivity to objective posterior probability: Subjective posterior probability must have been significantly higher (*α* = 0.05, two-sample t-test assuming unknown and unequal variances) on trials with a high objective posterior probability (Pr(*Q*|*x*) ≥ 0.9) than on trials with a low objective posterior probability (Pr(*Q*|*x*) ≤ 0.1).

To measure participants’ intrinsic posterior-estimation strategies without extensive training, the criteria were designed to be lenient enough to respect variation in their pre-task strategies while excluding participants who disengaged from the task or who adopted strategies clearly consistent with misunderstanding the task.

#### Earnings

Compensation for the prescan session was a show-up fee of $15 on top of their earnings from the payout trial (up to $30) on the prescan session. Compensation for the scan session was a show-up fee of $20 on top of their earnings from the payout trial (up to $30) on the scan session. Participants received an extra $50 for completing both sessions. Therefore, they could earn up to $145 for completing the entire study.

### Modeling Subjective Posterior Probability

From the participants’ perspective, the objective of the Probability Estimation Task is to maximize earnings by estimating the posterior probability of the questioned gallery conditional on the sample from the hidden gallery (Pr(*Q*|*x*)). According to Bayes’ theorem, this posterior probability is a function of the prior probability of the questioned gallery (Pr(*Q*)) and the likelihood of the sample conditional on the questioned gallery (Pr(*x*|*Q*)) (**Equation 5**).

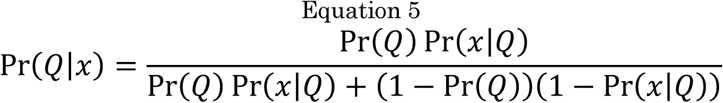

On each trial, the prior probability of the portrait and landscape galleries was explicitly stated while the likelihood was conveyed by the revealed sample picture and the sample’s evidence strength, displayed as the ratio of majority-category to minority-category pictures (i.e., 60:40; **Figure 1A**). For the purposes of the formulae, this ratio was converted into a probability with domain (0.5, 1] (e.g., 60:40 became 0.6). (However, all evidence strengths on the task were less than 1.) The relationship between evidence strength *θ* and likelihood Pr(*x*|*Q*)depended on the trial’s questioned gallery: Pr(*x*|*Q*) = *θ* when the sample signaled the questioned gallery (i.e., when the sample was a face and the questioned gallery was the portrait gallery, or when the sample was a place and the questioned gallery was the landscape gallery), and Pr(*x*|*Q*) = 1− *θ* when the sample did not signal the questioned gallery.

To measure the effects of prior probability and likelihood on participants’ reported subjective posteriors, we parameterized Bayes’ theorem. To start, we reexpressed it as the sum of log odds (logits), which converts the domain and range from the interval [0,1] to the interval (−∞, +∞) and allows us to model subjective posterior using linear regression (**Equation 6**, by convention, all logarithms in this paper use the natural logarithm).

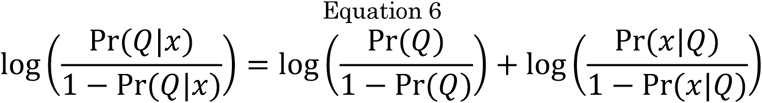

**Equation 6** simply states that the log posterior odds (logit posterior) is the sum of the log prior odds (logit prior) and the log likelihood odds (log likelihood ratio, logit likelihood). From here, we parameterized the influence of prior and likelihood on the subjective posterior probability (*π*(*Q*|*x*)) using mixed-effects regression.

To model subjective posterior probability as a function of the objective posterior probability, we included fixed-effects terms for the intercept and objective logit posterior along with the corresponding random-effects terms by participant (**Equation 7**).

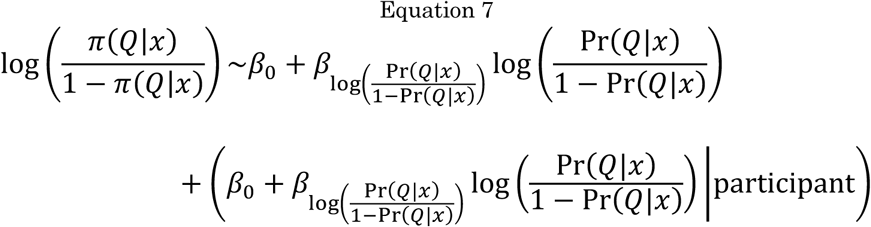

To model subjective posterior probability as a function of prior and likelihood, we included fixed- and random-effects terms for the intercept, logit prior, and logit likelihood (**Equation 8**).

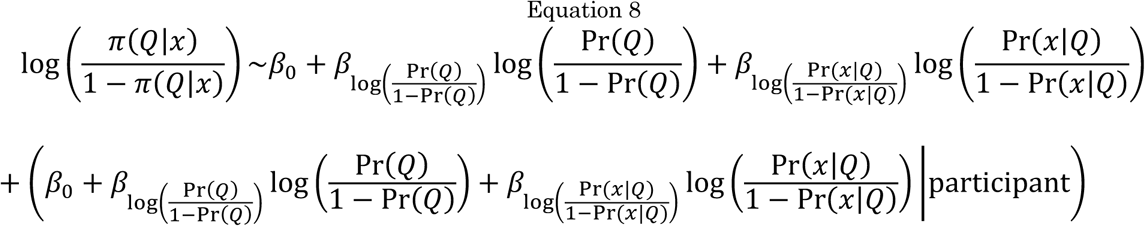

To account for the potentially confounding effects of penalty (*W*) and initial slider position (*S*), we added nuisance regressors for those terms to **Equation 8** (**Equation 9**).

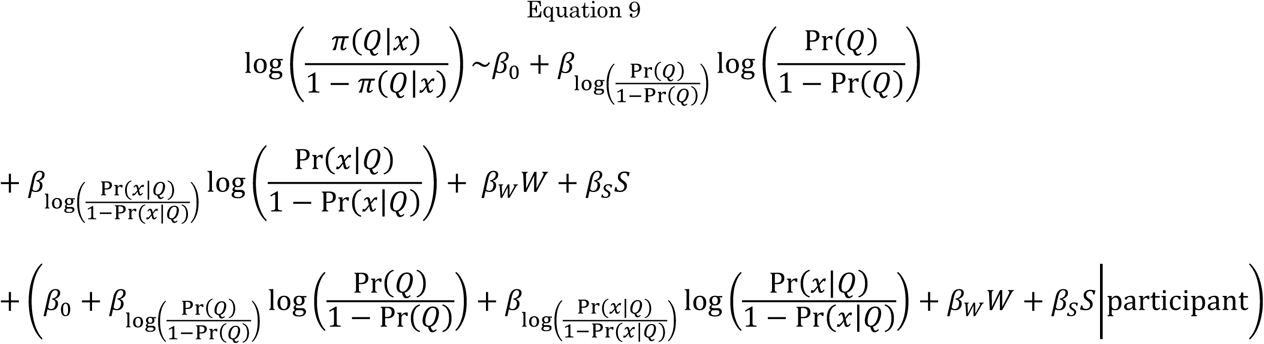

Model criterion scores for all tested models are plotted in **Figure S3**. Fixed-effects model coefficients (weights) are displayed in **Figure 2B, Figure S4**, and **Table S3– Table S5**. Weights were fit using maximum likelihood estimation. Because participants could only submit a subjective posterior probability between 0.02 and 0.98, inclusive, there was no risk that subjective logit posterior would equal −∞ or ∞, which would make the models inestimable. A corresponding “ideal” observer was simulated for each participant, submitting a posterior probability “estimate” as close to the objective posterior probability as possible within the limitations of the accepted responses on the slider.

### fMRI Data Acquisition and Preprocessing

Whole-brain fMRI data were acquired on a 3-T Siemens MAGNETOM Prisma scanner with a 64-channel head coil at the Magnetic Resonance Imaging Center at the Zuckerman Mind Brain Behavior Institute of Columbia University. Functional images were acquired with a T2*-weighted, two-dimensional gradient echo spiral in/out pulse sequence (repetition time (TR) = 1,000 ms; echo time = 30 ms; flip angle = 52°, field of view = 230 mm; 2.4×2.4×2.4 mm voxels; 56 slices; multiband factor = 4). To reduce dropout in central frontal regions, slices were tilted by 10° forward from the AC-PC axis. During the scan session, the behavioral tasks were projected onto a mirror attached to the scanner head coil for the participant to see (Hyperion MRI Digital Projection System); participants made responses with the right hand through an MRI-compatible trackball (Current Design). fMRI data were preprocessed using fMRIPrep. For details on procedures on registration and normalization of data, please see the Supplement.

### fMRI Data Analysis

Statistical analyses were conducted using the general linear model (GLM) framework implemented in SPM12, Version 7487 (https://www.fil.ion.ucl.ac.uk/spm), convolving boxcar functions and parametric modulators within the GLM by the SPM canonical hemodynamic response function. Statistical maps from functional data were overlaid on an average of the 23 participants’ individual T1-weighted (T1w) maps normalized to Montreal Neurological Institute (MNI) space and subsequently smoothed with a Gaussian kernel with a field-width at half-maximum (FWHM) of 5×5×5 mm to mimic the smoothness of the functional images used in the localization analyses. Since scanning did not occur during the Payout Stage, fMRI activation was only measured during the Estimation Stage.

#### Probability Estimation Task

##### Whole-brain localization analyses

Functional images normalized to MNI space were smoothed with a Gaussian kernel with a FWHM of 5×5×5 mm before a whole-brain localization analysis was performed with a summary statistics approach. First, a voxel-wise contrast map was estimated in a first-level (participant-level) analysis for every participant from their functional image time series. All first-level GLMs for whole-brain localization of predictors used a variable-epoch model to model participants’ responses^54^: each GLM contained one boxcar function to model the decision period (the period between the beginning of the response window and the reaction time on non-catch trials that received a response (**Figure 1A**; henceforth called the *response boxcar*) and another boxcar function to model the same period during catch trials. The response boxcar was parametrically modulated by a set of predictors that varied by localization GLM (see below). If the participant failed to respond to at least one trial during a run of the Estimation Stage, a third boxcar function was added to model the entire response window on the trials that they omitted. The localization GLMs also contained fixed-body motion-realignment regressors (*x, y, z*, pitch, roll, and yaw) and their respective first derivatives. Each participant’s contrast map was submitted to a second-level t-test at the group level, applying a cluster-wise correction for multiple comparison using non-parametric permutation tests in SnPM13.1.08 (http://nisox.org/Software/SnPM13/)^55^. Permutation tests were based on a stringent cluster-forming threshold of *p* = 0.001 and considered significant at a cluster-wise familywise error rate threshold of *p* < 0.05; we used 10,000 permutations and applied variance smoothing of group-level images by a Gaussian kernel with a FWHM of 5×5×5 mm, consistent with recommendations^55,56^.

In the GLM to localize subjective logit posterior according to participants’ reports (WB-GLM 1), the response boxcar was parametrically modulated by (1) the absolute value of the subjective logit posterior (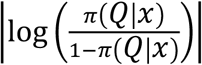, a representation of the subjective posterior certainty), (2) a dummy variable indicating if the subjective posterior favored the questioned gallery (yes = 1, no = −1), (3) a dummy variable indicating the side of the screen on which the sample picture appeared (sample on right = 1, sample on left = −1), (4) the penalty, (5) the subjective logit posterior of the questioned gallery (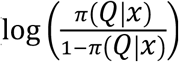, the product of modulators 1 and 2), (6) the subjective expected value (**Equation 4**), and (7) the initial slider position normalized to between 0 and 1.

In the GLM to localize *objective* logit posterior (WB-GLM 2), the response boxcar function was parametrically modulated by (1) the absolute value of the objective logit posterior (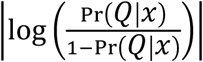, a representation of the objective posterior certainty), (2) a dummy variable indicating if the objective posterior favored the questioned gallery (yes = 1, no = −1), (3) a dummy variable indicating the side of the screen on which the sample picture appeared (sample on right = 1, sample on left = −1), (4) the penalty, (5) the objective logit posterior of the questioned gallery (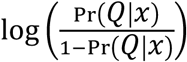, the product of modulators 1 and 2), (6) the objective expected value (**Equation 3**), and (7) the initial slider position normalized to between 0 and 1.

In the GLM to localize logit prior and logit likelihood (WB-GLM 3), the response boxcar was parametrically modulated by (1) the absolute value of the logit prior (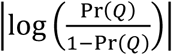, a representation of the prior certainty), (2) the absolute value of the logit likelihood (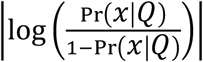, a representation of the likelihood certainty), (3) a dummy variable indicating if the logit prior favored the questioned gallery (yes = 1, no = −1), (4) a dummy variable indicating if the logit likelihood favored the questioned gallery (yes = 1, no = –1), (5) a dummy variable indicating the side of the screen on which the sample picture appeared (sample on right = 1, sample on left = −1), (6) the penalty, (7) the logit prior of the questioned gallery (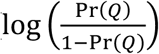, the product of modulators 1 and 3), (8) the logit likelihood conditional on the questioned gallery (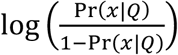, the product of modulators 2 and 4), (9) the subjective expected value (**Equation 4**), and (10) the initial slider position normalized to between 0 and 1.

We also designed a GLM (WB-GLM 4) to localize the model-fitted subjective logit posterior 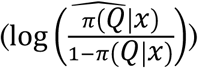 as estimated by **Equation 9**. Its response boxcar was parametrically modulated by (1) the absolute value of the model-fitted subjective logit posterior 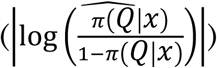, (2) a dummy variable indicating if the model-fitted subjective posterior favored the questioned gallery (yes = 1, no = −1), (3) a dummy variable indicating the side of the screen on which the sample picture appeared (sample on right = 1, sample on left = −1), (4) the penalty, (5) the model-fitted subjective logit posterior of the questioned gallery (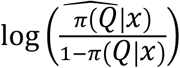, the product of modulators 1 and 2), (6) the subjective expected value (**Equation 4**, but replacing the subjective posterior probability with is model-fitted version), and (7) the initial slider position normalized to between 0 and 1. SPM orthogonalization was turned off for the first-level analyses. We assessed the contrast between the effects of logit prior (of the questioned gallery) and logit likelihood (of the sample, conditional on the questioned gallery) (**Figure S8**) by using a conjunction null, defined as regions that showed significant activation by both effects ^57^. The fact that all the trials within a run had the same questioned gallery allowed us to apply contrasts to the modulators for the subjective logit posterior, logit prior, and logit likelihood to create contrasts maps of these logits with respect to the questioned gallery, portrait gallery, or landscape gallery.

##### fROI analyses

We used a functional region of interest (fROI) approach to measure average activation tracking different predictors within a cluster or region of interest^58^. We did so by defining first-level GLMs to create whole-brain contrast maps showing the response of each voxel to a particular predictor of interest. All these GLMs were fit to participants’ normalized but unsmoothed functional time series from the Probability Estimation Task except for the analyses of the face- and place-selective fROIs, which were fit to functional time series that were registered to participants’ individual, unnormalized T1w images. Then, we measured the average contrast statistic of that variable within the fROI of interest. These fROI GLMs consisted of the same boxcar functions and motion regressors as the localization GLMs but varied by the parametric modulators for the response boxcar function.

In the GLM to measure activation tracking prior and likelihood (fROI-GLM 1), the response boxcar was parametrically modulated by (1) the logit prior of the questioned gallery (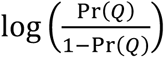), (2) the logit likelihood conditional on the questioned gallery (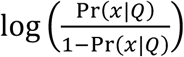), (3) penalty, and (4) slider displacement (the difference between the initial slider position and the slider position at submission of the response). In the GLM to measure activation by the objective logit posterior (fROI-GLM 2), the response boxcar was parametrically modulated by (1) the objective logit posterior 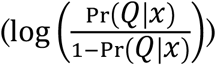, (2) penalty, and (3) slider displacement. In the GLM to measure activation by the quadratic term for subjective logit posterior (fROI-GLM 3), the response boxcar was parametrically modulated by (1) the subjective logit posterior of the questioned gallery 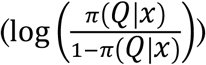, (2) the square of this term 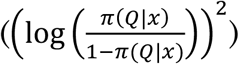, (3) penalty, and (4) slider displacement. SPM orthogonalization was turned off, and appropriate contrasts were applied to the logit predictors to create contrast maps of these logits with respct to the questioned gallery, portrait gallery, or landscape gallery.

Note that the parametric modulator for *slider displacement* in the fROI GLMs is different from the parametric modulator for *initial slider position* in the localization GLMs. While initial slider position was included in the localization GLMs to control for eye and hand movement, we did not include *slider displacement* in those GLMs because slider displacement is correlated with the participant’s response of interest: the subjective logit posterior. In the localization GLMs, we did not want the predictors to account for variance attributable to the participant’s response, while in the fROI GLMs, we wanted to introduce a more stringent test to determine if the variables of interest could survive the nuisance confound.

We used mixed-effects regression to test the effect of the predictors in each fROI analysis except the analysis to test differential activation of the face- and place-selective fROIs sorted by questioned gallery (see later). For each mixed-effects regression test, the dependent variable was a vector of all contrast statistics for each voxel from each relevant fROI from each participant. In analyses of only one fROI, the independent variables were dummy variables indicating the contrast map from which the corresponding contrast statistic was derived, with random effects by participant. In analyses that included more than one fROI, the independent variables were dummy variables indicating the combination of fROI and contrast map from which the corresponding contrast statistic was derived, with random effects by participant. We used the fixed-effects coefficients from these analyses as representations of the average effect of each predictor of interest.

To test the average effect of logit prior and logit likelihood within the PPC cluster (**Figure 3C**; **Figure S6B–C**), we estimated voxel-wise contrast statistics from the PPC cluster by using fROI-GLM 1 and modeled them as a function of the contrast from which they had been derived (**Equation 10**).

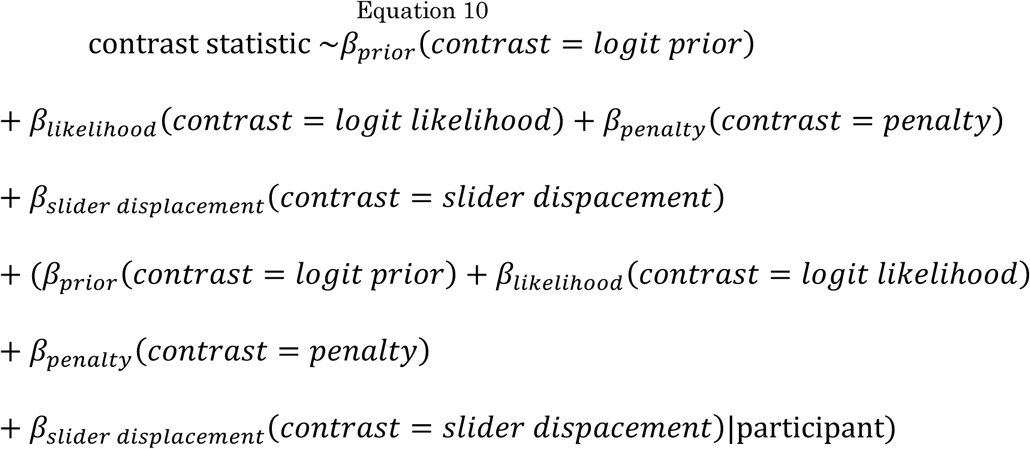

To test the average effect of the quadratic term for subjective logit posterior within the PPC cluster (**Table S7**), we estimated voxel-wise contrast statistics from the PPC cluster by using fROI-GLM 3 and modeled them as a function of the contrast from which they had been derived (**Equation 11**).

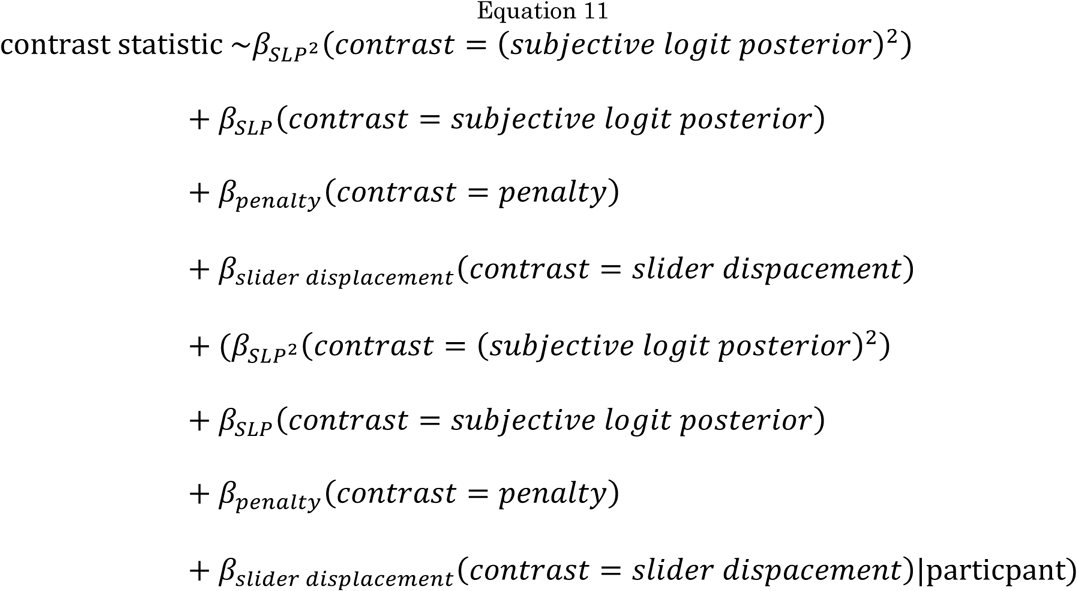

To test the average effect of the objective logit posterior within the PPC cluster (**Figure 3E, Figure S6A**), we estimated voxel-wise contrast statistics from the PPC cluster by using fROI-GLM 4 and modeled them as a function of the contrast maps from which they had been derived (**Equation 12**).

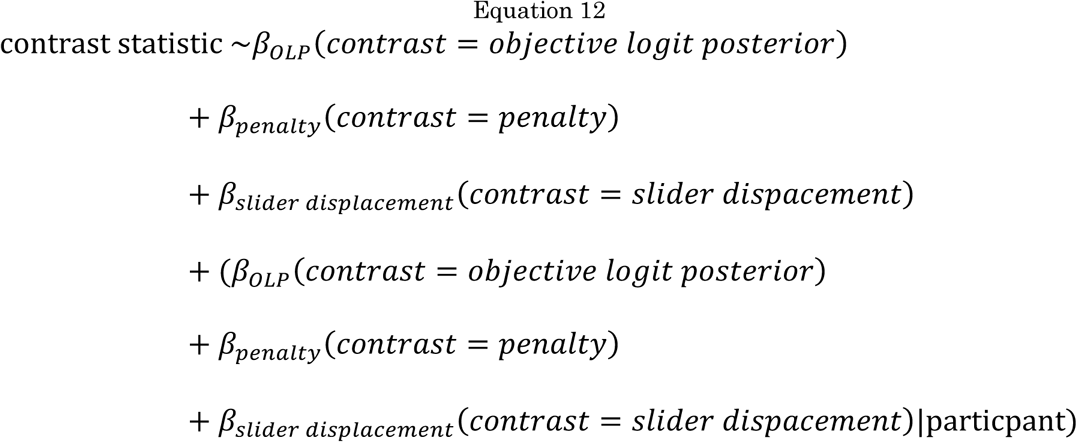

To test the average effect of subjective logit posterior within face- and place-selective regions (**Figure 4B, Figure S9A**), we estimated voxel-wise contrast statistics from every participant’s face and place fROI by using fROI-GLM 2 applying contrasts to convert the reference frame of the logits from the questioned gallery to the portrait and landscape galleries) and modeled them as a function of the contrasts and the fROI (face- or place-selective) from which they had been derived (**Equation 13**).

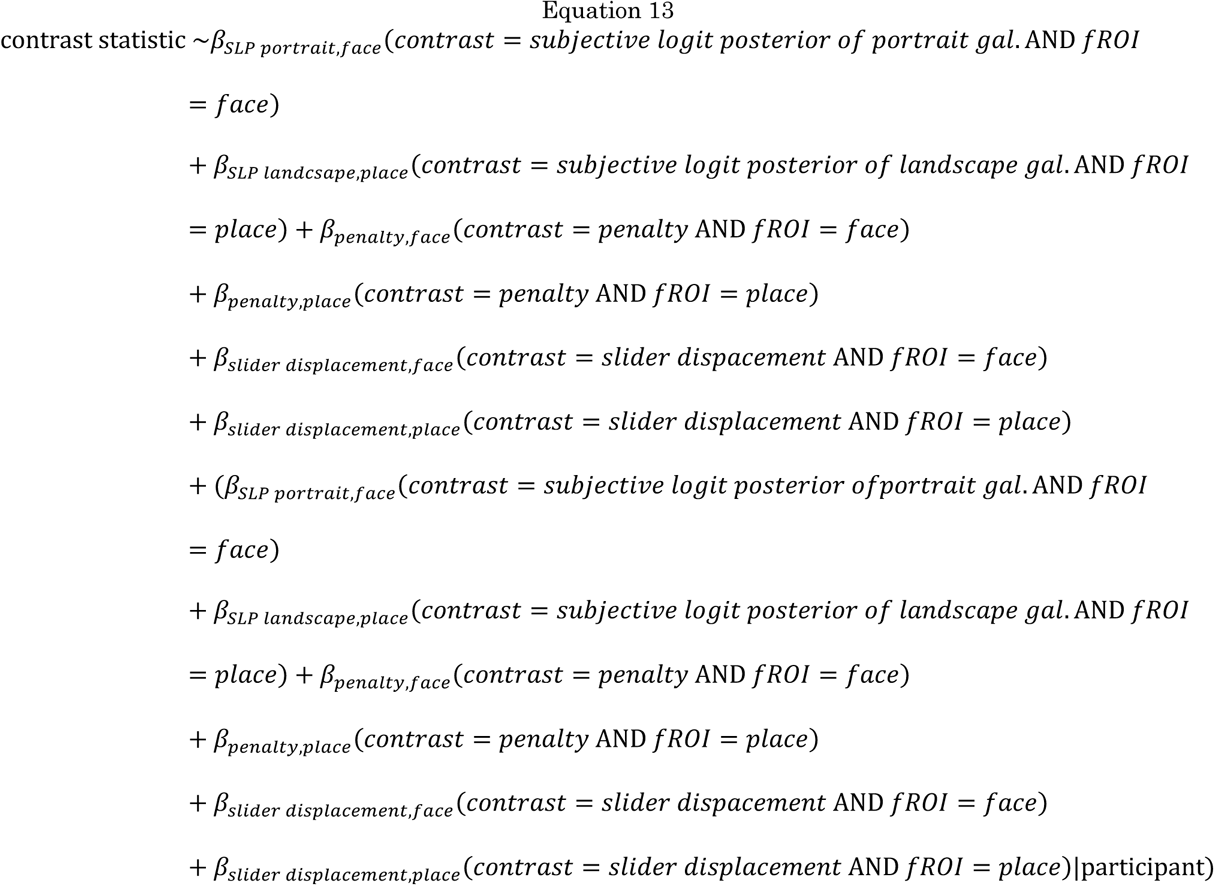

To test the average effect of logit prior and logit likelihood within face- and place-selective regions (**Figure S9B–C**), we estimated voxel-wise contrast statistics from every participant’s face and place fROI by using fROI-GLM 1 (applying contrasts to convert the reference frame of the logits from the questioned gallery to the portrait and landscape galleries) and modeled them as a function of the contrasts and the fROI (face- or place-selective) from which they had been derived (**Equation 14**).

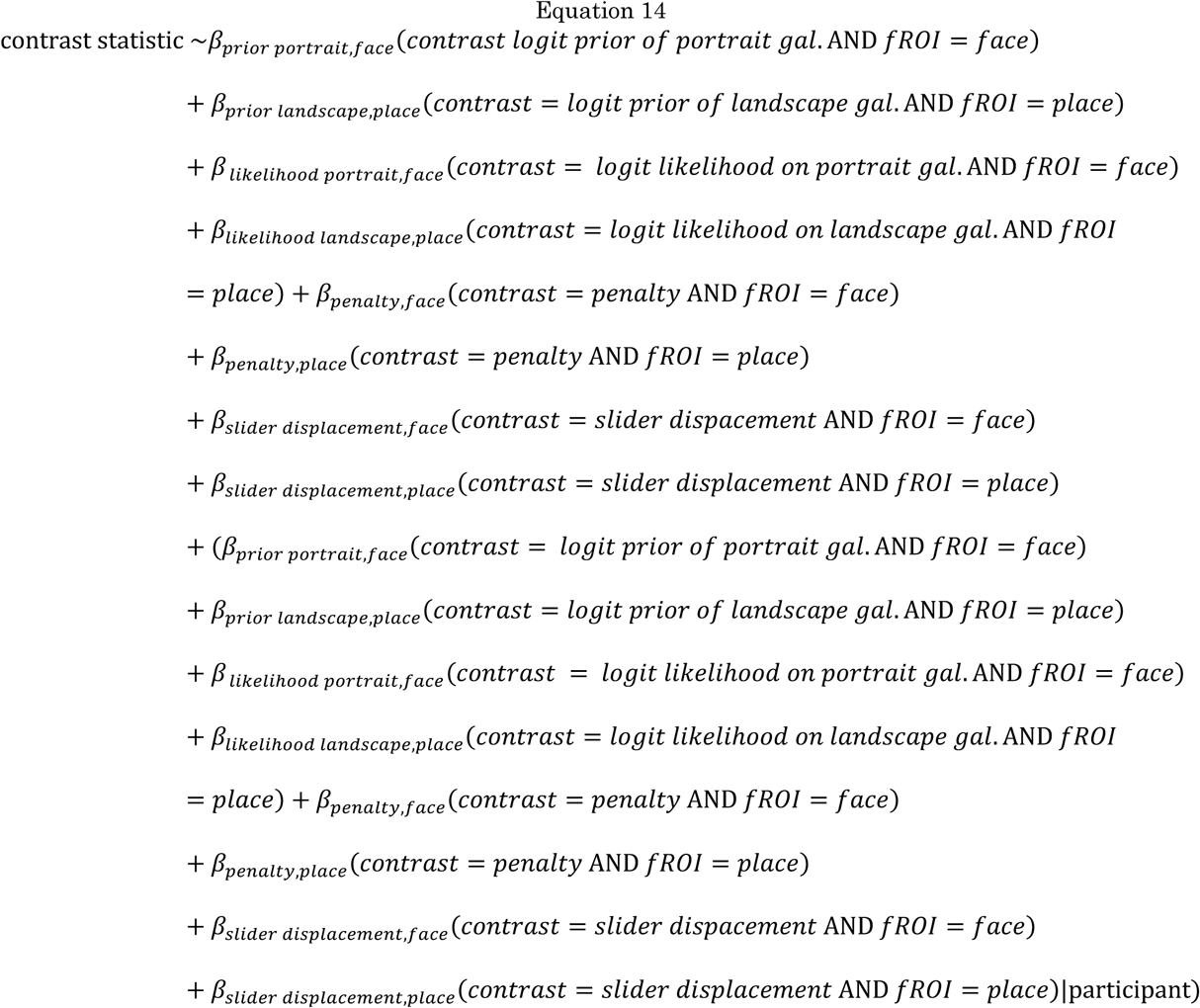

When sorting trials by the questioned gallery, we used a summary statistics approach to test the differential effect of subjective logit posterior, logit prior, and logit likelihood within the face- and place-selective fROIs. In this analysis, we only examined logits with respect to the questioned gallery (e.g., the subjective logit posterior of the portrait gallery only during runs where the portrait gallery was the questioned gallery) to account for framing effects created by the questioned gallery. To do so, we calculated the mean contrast statistic in each participant’s face- and place-selective fROI, and we used an ANOVA to test for a significant interaction between the fROI and the contrast statistics for subjective logit posterior, logit prior, and logit likelihoods with respect to the gallery categories.

##### Face-Place Localizer

We defined a GLM to localize face- and place-selective regions of occipital and temporal cortex during the Face-Place Localizer Task in each participant. Unlike the GLMs for belief-updating localization and verification, the face-place localizer analysis was applied to functional volumes in each participant’s native brain space (without normalization). The first-level GLM consisted of two boxcar functions: one representing the period during which a face picture appeared on screen and another representing the period during which a place picture appeared on screen. Boxcar functions were convolved with the SPM canonical hemodynamic response function. Localization analysis was small volume–corrected to include only the occipital and temporal lobes in each participant. Face-selective regions were defined with the contrast “Face > Place,” while place-selective regions were defined with the contrast “Place > Face,” both at an uncorrected *p*-value threshold of 0.001 at the individual level.

## Acknowledgements

We would like to thank Chuen-Shin (Jessica) Cheng, Dania Elder, and Laura Hunter for help with data collection; Greg Jensen for help with data analysis; Kenneth Wengler for help with MRI scanner sequences; and Mariam Aly, Kenneth Wengler, Daniel Wolpert, and Michael Woodford for helpful advice on the project and manuscript. This study was funded by the Seed Grant for MR Studies Program of the Zuckerman Mind Brain Behavior Institute at Columbia University (CU-ZI-MR-S-0011) and the National Science Foundation Graduate Research Fellowship Program (DGE-1644869, NMS). This work was also supported by the National Institute of Mental Health under awards R01MH117323 and R01MH114965 (GH).

## References

1. Hemsley, D. R. & Garety, P. A. The Formation of Maintenance of Delusions: a Bayesian Analysis. Br. J. Psychiatry 149, 51–56 (1986).

2. Fletcher, P. C. & Frith, C. D. Perceiving is believing: A Bayesian approach to explaining the positive symptoms of schizophrenia. Nat. Rev. Neurosci. 10, 48–58 (2009).

3. Coltheart, M., Menzies, P. & Sutton, J. Abductive inference and delusional belief. http://dx.doi.org/10.1080/13546800903439120 15, 261–287 (2009).

4. Adams, R. A., Stephan, K. E., Brown, H. R., Frith, C. D. & Friston, K. J. The computational anatomy of psychosis. Front. Psychiatry 4, 47 (2013).

5. Fischhoff, B. & Beyth-Marom, R. Hypothesis evaluation from a Bayesian perspective. Psychol. Rev. 90, 239–260 (1983).

6. El-Gamal, M. A. & Grether, D. M. Are people Bayesian? Uncovering behavioral strategies. J. Am. Stat. Assoc. 90, 1137–1145 (1995).

7. Coutts, A. Good news and bad news are still news: experimental evidence on belief updating. Exp. Econ. 2018 222 22, 369–395 (2018).

8. Benjamin, D. J. Errors in probabilistic reasoning and judgment biases. in Handbook in Behavioral Economics, Volume 2 69–186 (2019). doi:10.1016/bs.hesbe.2018.11.002

9. Forstmann, B. U., Brown, S., Dutilh, G., Neumann, J. & Wagenmakers, E. J. The neural substrate of prior information in perceptual decision making: A model-based analysis. Front. Hum. Neurosci. 4, 1–12 (2010).

10. FitzGerald, T. H. B., Seymour, B., Bach, D. R. & Dolan, R. J. Differentiable neural substrates for learned and described value and risk. Curr. Biol. 20, 1823–1829 (2010).

11. Philiastides, M. G., Biele, G. & Heekeren, H. R. A mechanistic account of value computation in the human brain. Proc. Natl. Acad. Sci. 107, 9430–9435 (2010).

12. d’Acremont, M., Fornari, E. & Bossaerts, P. Activity in Inferior Parietal and Medial Prefrontal Cortex Signals the Accumulation of Evidence in a Probability Learning Task. PLoS Comput. Biol. 9, 1002895 (2013).

13. d’Acremont, M., Schultz, W. & Bossaerts, P. The Human Brain Encodes Event Frequencies While Forming Subjective Beliefs. J. Neurosci. 33, 10887–10897 (2013).

14. Mengotti, P., Dombert, P. L., Fink, G. R. & Vossel, S. Disruption of the right temporoparietal junction impairs probabilistic belief updating. J. Neurosci. 37, 5419–5428 (2017).

15. Machina, M. J. & Schmeidler, D. A More Robust Definition of Subjective Probability. Econometrica 60, 745 (1992).

16. Charpentier, C. J., Bromberg-Martin, E. S. & Sharot, T. Valuation of knowledge and ignorance in mesolimbic reward circuitry. Proc. Natl. Acad. Sci. U. S. A. 115, E7255–E7264 (2018).

17. Kobayashi, K., Ravaioli, S., Baranès, A., Woodford, M. & Gottlieb, J. Diverse motives for human curiosity. Nat. Hum. Behav. 3, 587–595 (2019).

18. Ting, C. C. et al. Neural Mechanisms for Integrating Prior Knowledge and Likelihood in Value-Based Probabilistic Inference. J. Neurosci. 35, 1792–1805 (2015).

19. Huk, A. C. & Shadlen, M. N. Neural activity in macaque parietal cortex reflects temporal integration of visual motion signals during perceptual decision making. J. Neurosci. 25, 10420–10436 (2005).

20. Kanwisher, N., McDermott, J. & Chun, M. M. The Fusiform Face Area: A Module in Human Extrastriate Cortex Specialized for Face Perception. J. Neurosci. 17, 4302–4311 (1997).

21. Epstein, R. & Kanwisher, N. A cortical representation of the local visual environment. Nature 392, 598–601 (1998).

22. Gold, J. I. & Shadlen, M. N. Neural computations that underlie decisions about sensory stimuli. Trends in Cognitive Sciences (2001). doi:10.1016/S1364-6613(00)01567-9

23. Ditterich, J. Stochastic models of decisions about motion direction: Behavior and physiology. Neural Networks 19, 981–1012 (2006).

24. Carpenter, R. H. S. & Williams, M. L. L. Neural computation of log likelihood in control of saccadic eye movements. Nat. 1995 3776544 377, 59–62 (1995).

25. Roitman, J. D. & Shadlen, M. N. Response of Neurons in the Lateral Intraparietal Area during a Combined Visual Discrimination Reaction Time Task. J. Neurosci. 22, 9475–9489 (2002).

26. Zhang, H. & Maloney, L. T. Ubiquitous log odds: A common representation of probability and frequency distortion in perception, action, and cognition. Front. Neurosci. 6, (2012).

27. Garcia, B., Cerrotti, F. & Palminteri, S. The description-experience gap: A challenge for the neuroeconomics of decision-making under uncertainty. Philos. Trans. R. Soc. B Biol. Sci. 376, (2021).

28. Hanks, T. D., Ditterich, J. & Shadlen, M. N. Microstimulation of macaque area LIP affects decision-making in a motion discrimination task. Nat. Neurosci. 9, 682–689 (2006).

29. Culham, J. C. & Kanwisher, N. G. Neuroimaging of cognitive functions in human parietal cortex. Curr. Opin. Neurobiol. 11, 157–163 (2001).

30. Bisley, J. W. & Goldberg, M. E. Attention, Intention, and Priority in the Parietal Lobe. Annu. Rev. Neurosci. 33, 1–21 (2010).

31. Platt, M. L. & Glimcher, P. W. Neural correlates of decision variables in parietal cortex. Nature 400, 233–238 (1999).

32. Rutledge, R. B., Dean, M., Caplin, A. & Glimcher, P. W. Testing the Reward Prediction Error Hypothesis with an Axiomatic Model. J. Neurosci. 30, 13525– 13536 (2010).

33. Roy, M. et al. Representation of aversive prediction errors in the human periaqueductal gray. Nat. Neurosci. 17, 1607–1612 (2014).

34. Sereno, M. I., Pitzalis, S. & Martinez, A. Mapping of contralateral space in retinotopic coordinates by a parietal cortical area in humans. Science (80-.). 294, 1350–1354 (2001).

35. Grefkes, C. & Fink, G. R. The functional organization of the intraparietal sulcus in humans and monkeys. Journal of Anatomy 207, 3–17 (2005).

36. Glasser, M. F. & Van Essen, D. C. Mapping human cortical areas in vivo based on myelin content as revealed by T1- and T2-weighted MRI. J. Neurosci. 31, 11597–616 (2011).

37. Kira, S., Yang, T. & Shadlen, M. N. A Neural Implementation of Wald’s Sequential Probability Ratio Test. Neuron 85, 861–873 (2015).

38. Foley, N. C., Kelly, S. P., Mhatre, H., Lopes, M. & Gottlieb, J. Parietal neurons encode expected gains in instrumental information. Proc. Natl. Acad. Sci. 114, E3315–E3323 (2017).

39. Horan, M., Daddaoua, N. & Gottlieb, J. Parietal neurons encode information sampling based on decision uncertainty. Nat. Neurosci. 22, 1327–1335 (2019).

40. Li, Y., Daddaoua, N., Horan, M. & Gottlieb, J. Uncertainty modulates visual maps during non-instrumental information demand. bioRxiv 2021.07.20.453107 (2021). doi:10.1101/2021.07.20.453107

41. Wisniewski, D., Reverberi, C., Momennejad, I., Kahnt, T. & Haynes, J.-D. The Role of the Parietal Cortex in the Representation of Task–Reward Associations. J. Neurosci. 35, 12355–12365 (2015).

42. Beck, J. M. et al. Probabilistic Population Codes for Bayesian Decision Making. Neuron 60, 1142–1152 (2008).

43. Yang, T. & Shadlen, M. N. Probabilistic reasoning by neurons. Nature 447, 1075–1080 (2007).

44. Soltani, A. & Wang, X. J. Synaptic computation underlying probabilistic inference. Nat. Neurosci. 13, 112–119 (2010).

45. Simon, O., Mangin, J. F., Cohen, L., Le Bihan, D. & Dehaene, S. Topographical layout of hand, eye, calculation, and language-related areas in the human parietal lobe. Neuron 33, 475–487 (2002).

46. Ashkenazi, S., Rosenberg-Lee, M., Tenison, C. & Menon, V. Weak task-related modulation and stimulus representations during arithmetic problem solving in children with developmental dyscalculia. Dev. Cogn. Neurosci. 2, S152– S166 (2012).

47. Hertwig, R., Barron, G., Weber, E. U. & Erev, I. Decisions from experience and the effect of rare events in risky choice. Psychol. Sci. 15, 534–539 (2004).

48. Brainard, D. H. The Psychophysics Toolbox. Spat. Vis. 10, 433–436 (1997).

49. Kleiner, M., Brainard, D. & Pelli, D. What’s new in Psychtoolbox-3? Percept. ECVP Abstr. 36, (2007).

50. Righi, G., Peissig, J. J. & Tarr, M. J. Recognizing disguised faces. Vis. cogn. 20, 143–169 (2012).

51. Konkle, T., Brady, T. F., Alvarez, G. A. & Oliva, A. Scene memory is more detailed than you think: The role of categories in visual long-term memory. Psychol. Sci. 21, 1551–1556 (2010).

52. Hagberg, G. E., Zito, G., Patria, F. & Sanes, J. N. Improved detection of event-related functional MRI signals using probability functions. Neuroimage 14, 1193–1205 (2001).

53. Hossain, T. & Okui, R. The binarized scoring rule. Rev. Econ. Stud. 80, 984– 1001 (2013).

54. Grinband, J., Wager, T. D., Lindquist, M., Ferrera, V. P. & Hirsch, J. Detection of time-varying signals in event-related fMRI designs. Neuroimage 43, 509–520 (2008).

55. Nichols, T. E. & Holmes, A. P. Nonparametric permutation tests for functional neuroimaging: A primer with examples. Hum. Brain Mapp. 15, 1–25 (2002).

56. Holmes, A. P., Blair, R. C., Watson, J. D. G. & Ford, I. Nonparametric analysis of statistic images from functional mapping experiments. J. Cereb. Blood Flow Metab. 16, 7–22 (1996).

57. Nichols, T., Brett, M., Andersson, J., Wager, T. & Poline, J. B. Valid conjunction inference with the minimum statistic. Neuroimage 25, 653–660 (2005).

58. Saxe, R., Brett, M. & Kanwisher, N. Divide and conquer: A defense of functional localizers. Neuroimage 30, 1088–1096 (2006).

59. Eickhoff, S. B. et al. A new SPM toolbox for combining probabilistic cytoarchitectonic maps and functional imaging data. Neuroimage 25, 1325– 1335 (2005).

60. Eickhoff, S. B., Heim, S., Zilles, K. & Amunts, K. Testing anatomically specified hypotheses in functional imaging using cytoarchitectonic maps. Neuroimage 32, 570–582 (2006).

61. Eickhoff, S. B. et al. Assignment of functional activations to probabilistic cytoarchitectonic areas revisited. Neuroimage 36, 511–521 (2007).

62. Desikan, R. S. et al. An automated labeling system for subdividing the human cerebral cortex on MRI scans into gyral based regions of interest. Neuroimage 31, 968–980 (2006).

