## Supplement for "A neural substrate for Bayesian integration in human parietal cortex"

### Supplemental Figures

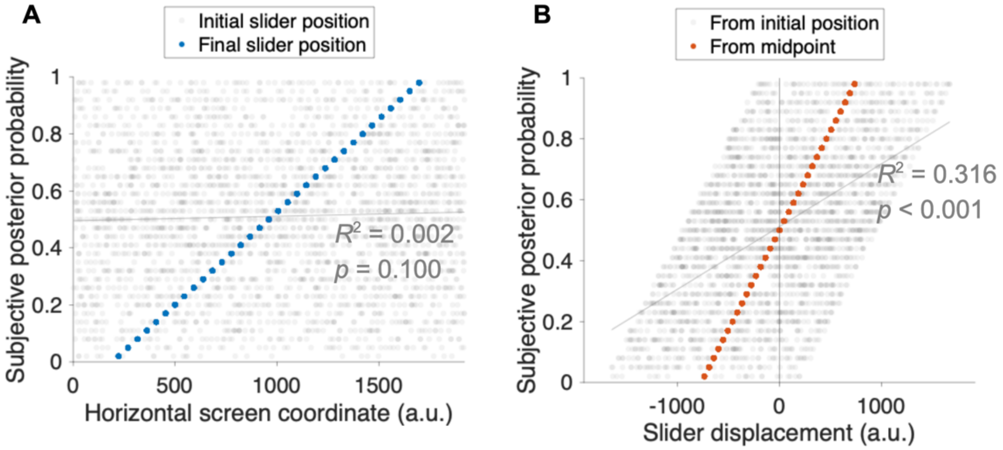


Figure S1. Randomizing the initial slider position to reduce its potential effect on the reported (subjective) posterior probability.

**A:** Translucent gray points represent the corresponding horizontal coordinate of the initial slider position ($x$-axis) for every reported (subjective) posterior probability ($y$-axis) in every non-catch trial from the scan session across all 23 participants. Regressing subjective posterior probability as a function of initial slider position with a mixed-effects model indicates no significant correlation between the two variables as determined by the model's coefficient of determination and the $p$-value of the fixed-effects coefficient for initial slider position ($R^2$ = 0.002, DF = 2,748, $p = 0.100$). Initial slider position was drawn from a random uniform distribution from the leftmost (0) to the rightmost (1920) horizontal screen coordinate in pixels. Translucent gray points represent initial slider position for all trials. Gray trendline is the least squares line for the relationship between subjective posterior probability and initial slider position. Solid blue points represent the centers of each posterior report bin on the slider (an estimate of the final slider position). The slider was divided into nonoverlapping posterior probability bins, increasing by steps of 3 percent (0.03) from 2 percent (0.02) on the left to 98 percent (0.98) on the right, making each bin approximately 46 pixels wide. Note that the slider could be placed to the left or right of the accepted responses.

**B:** Randomizing the initial slider position by trial reduces the correlation between the slider displacement and reported (subjective) posterior probability versus fixing the initial slider position to one point. Regressing subjective posterior probability as a function of slider displacement (see Methods) with a mixed-effects model indicates only a moderate positive correlation between the two variables as determined by the model's coefficient of determination and the $p$-value of the fixed-effects coefficient for slider displacement ($R^2$ = 0.316, DF = 2,748, $p < 0.001$); theoretically, this correlation would be almost perfect if the initial slider position had been stable across trials. Translucent gray points represent the corresponding estimated slider displacement ($x$-axis) for every subjective posterior probability ($y$-axis) in every non-catch trial from the scan session across all 23 participants. Gray diagonal line is the trendline for the subjective posterior probability as a function of slider displacement. The fact that the model's fixed-effects intercept is significantly greater than 0.5 counterintuitively indicates a slight *leftward* bias in participants' responses as a function of slider displacement alone (intercept: 0.512, SE = 0.004, DF = 2,748, $p = 0.003$ for difference from 0.5); intercept less than 0.5 would indicate a rightward bias). Estimated slider displacement was calculated as the difference between the mean slider position for the posterior report bin (an estimate of the final slider position) and the initial slider position, with negative differences representing leftward displacement and positive differences representing rightward displacement. Solid orange points represent the displacement of the center of each posterior probability bin from a fixed position on the slider (the midpoint).

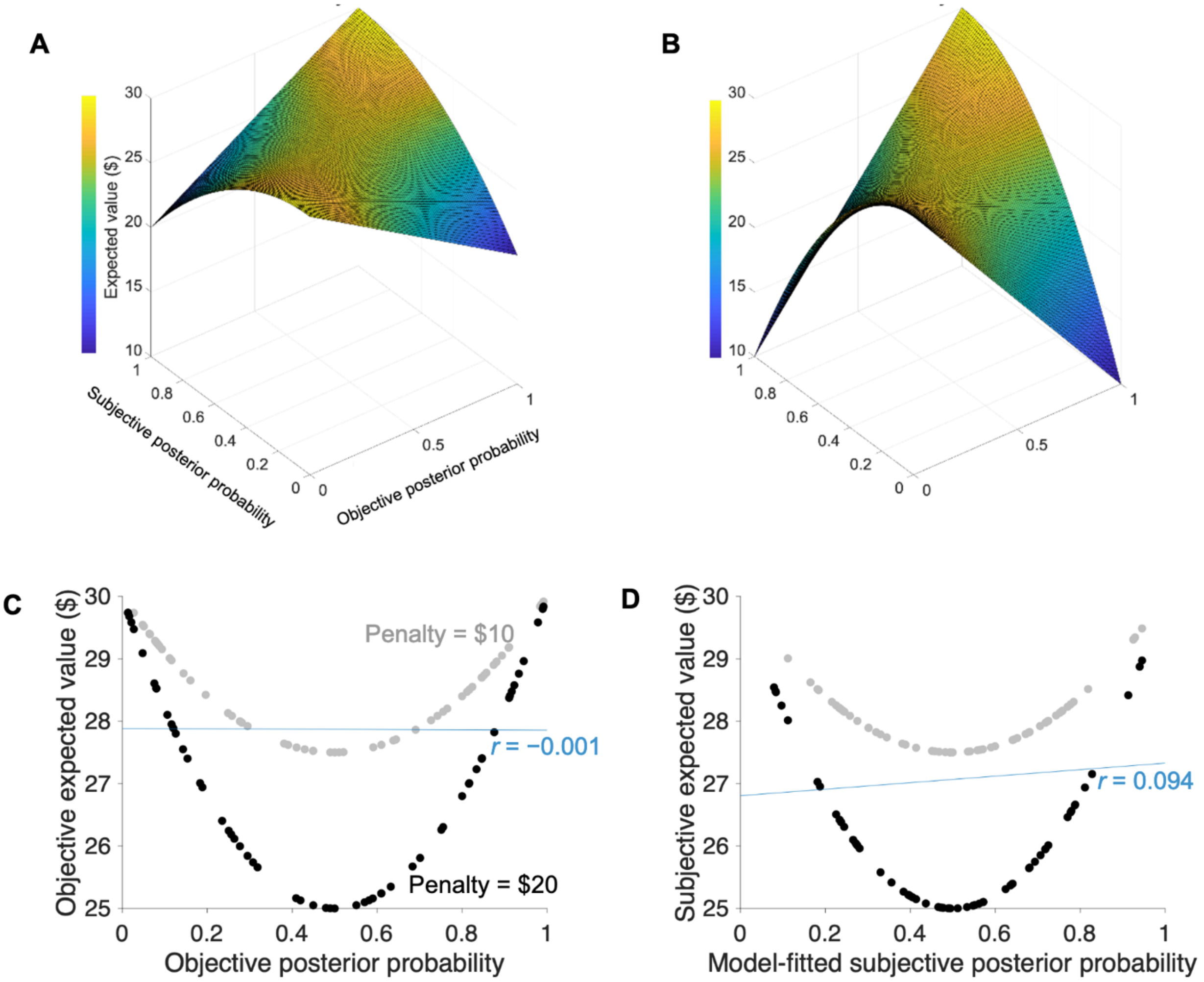


Figure S2. Expected value of a trial based on the binarized scoring rule with a quadratic cost function (see Methods).

**A:** Expected value of a trial in the $10 penalty condition over the entire space of possible subjective (submitted) and objective posterior probabilities.

**B:** Same as **A**, but for the $20 penalty condition.

**C:** In the imaginary situation that a participant submitted the exact objective posterior probability, there is only a negligible correlation between posterior probability and the expected value across all non-catch trials (Pearson correlation: −0.001, $N = 120$, no $p$-value because this is across the entire population of non-catch trials, not a sample of them).

**D:** There is a very low correlation between the model-fitted (**Equation 4**) subjective posterior probability and subjective expected value across all non-catch trials (Pearson correlation: 0.094, $N = 120$), assuming that participants were self-consistent and that they sought to minimize the cost function.

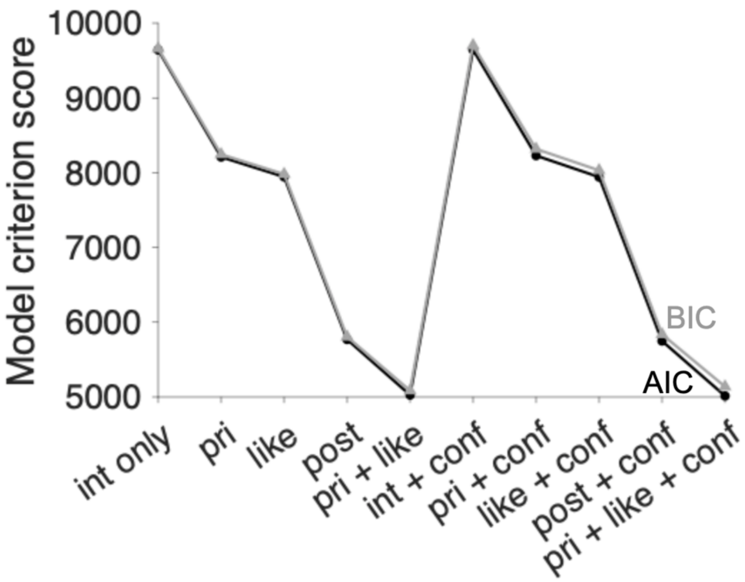


Figure S3. Akaike Information Criterion (AIC, black) and Bayesian Information Criterion (BIC, gray) scores for the various models of subjective logit posterior of the questioned gallery suggest that including separate regressors for logit prior and logit likelihood improves model accuracy compared to modeling subjective logit posterior only as a function of individual variables. Compares mixed-effects regression models that contain fixed- and random-effects terms for every regressor.

Regressor names:
"int only": intercept only
"pri": logit prior of questioned gallery
"like": logit likelihood of sample conditional on questioned gallery
"post": objective logit posterior of questioned gallery
"conf": penalty and initial slider position

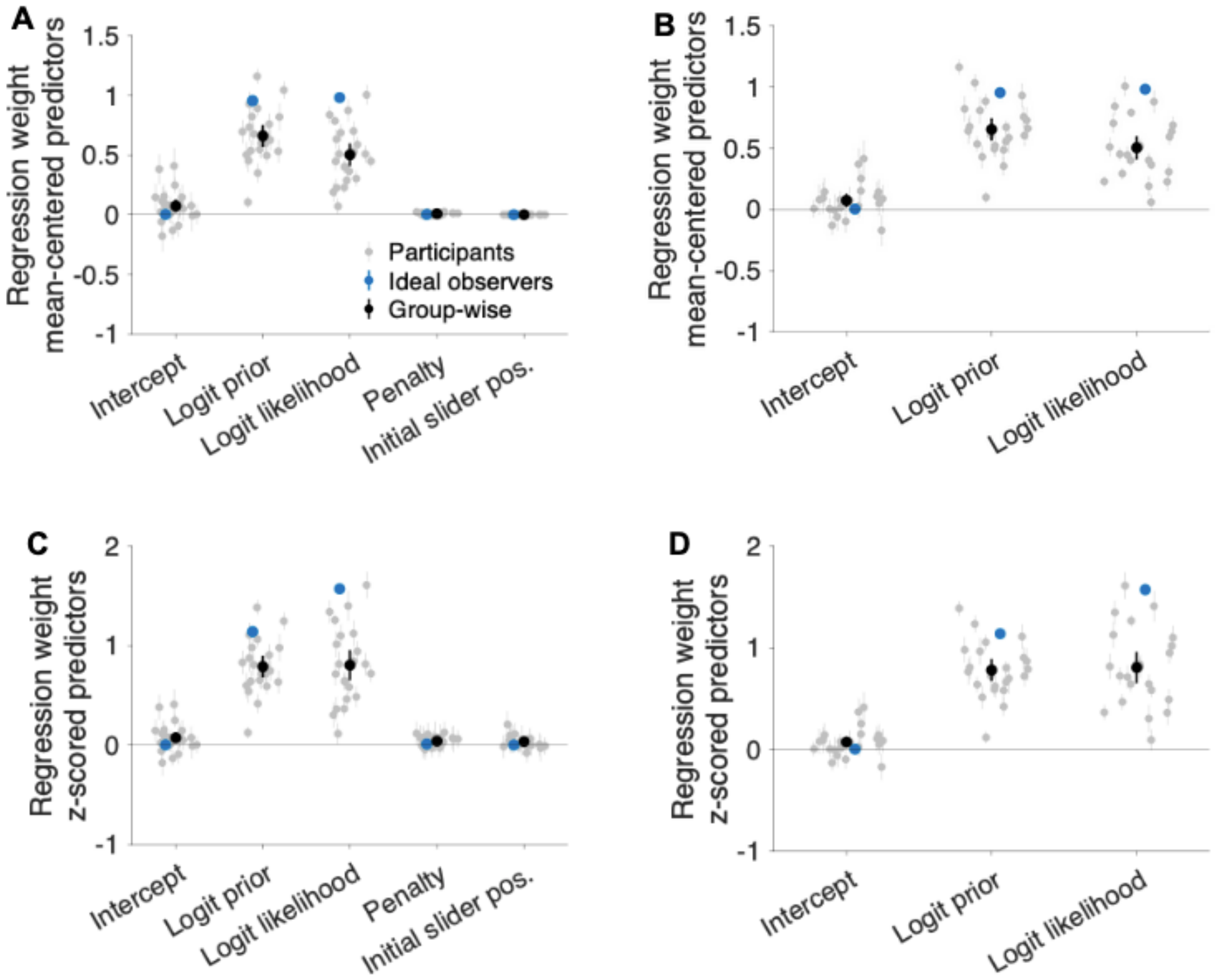


Figure S4. Participants integrate prior and likelihoods in their estimates of the subjective posterior of the questioned gallery, and the effect survives even after controlling for penalty and initial slider position. Error bars represent 95% confidence intervals.

**A:** Duplicate of **Figure 2C**. Group-level regression weights from the extended model of subjective logit posterior after mean-centering the predictors. Note that the logit prior and logit likelihood weights for the ideal observers are close to 1, as predicted by the implied coefficients in **Equation 6**.

**B:** Same as A, but for the reduced model of subjective logit posterior.

**C:** Same as A, but after z-scoring instead of simply mean-centering the predictors before fitting the model (**Table S5**).

**D:** Same as A, but for the reduced model of subjective logit posterior and after z-scoring the predictors before fitting the model (**Table S4**).

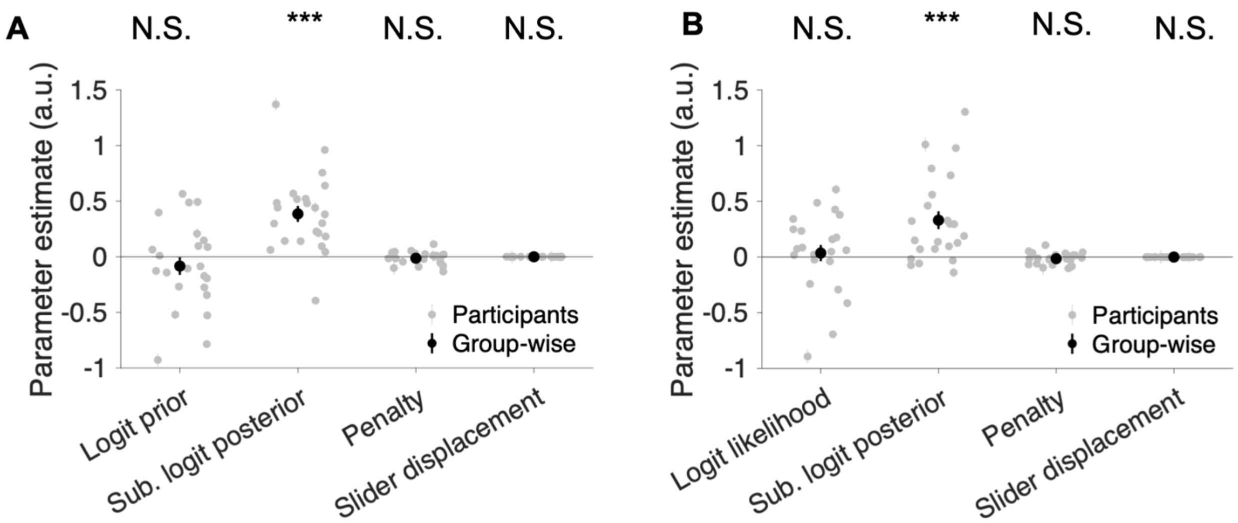


Figure S5. The effects of logit prior (**A**) and logit likelihood (**B**) within the posterior belief–encoding cluster (**Figure 3A**) do not remain statistically significant after accounting for subjective logit posterior, showing that they cannot be linearly dissociated from the subjective posterior signal within the cluster. ***: $p < 0.001$. N.S.: not significant (at $\alpha = 0.05$).

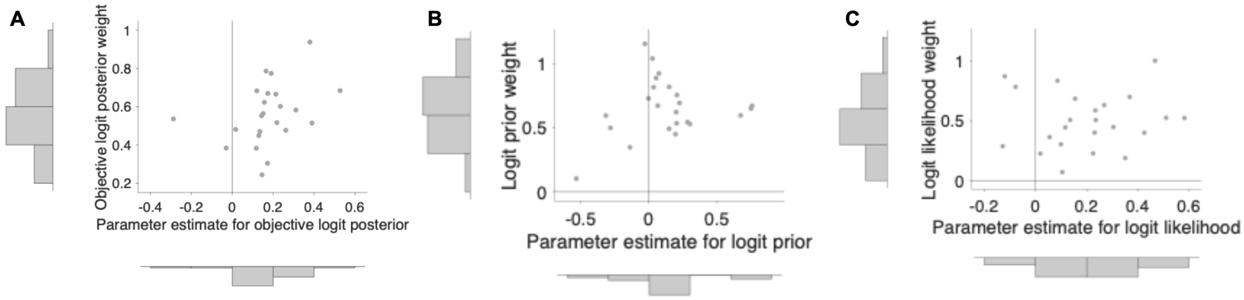


Figure S6. The sensitivity of the PPC cluster's signal to the objective posterior was positively correlated with behavioral sensitivity to objective logit posterior across all 23 participants, but signal sensitivity to prior and likelihood were not significantly correlated with their behavioral weights.

**A:** Duplicate of **Figure 3E**. BOLD signal to objective logit posterior (*x*-axis) within the PPC cluster is positively correlated with behavioral objective logit posterior weight (*y*-axis) across all participants (Spearman correlation: 0.439, $p = 0.037$), suggesting that distortions in neural representations of posterior probability in PPC contribute to the degree of distortion in participants' subjective posterior probabilities. Each point ($N = 23$) represents one participant.

**B:** BOLD signal to logit prior (*x*-axis) within the PPC cluster is not significantly correlated with behavioral logit prior weight (*y*-axis) (Spearman correlation: 0.008, $p = 0.973$). Each point ($N = 23$) represents one participant.

**C:** BOLD signal to logit likelihood (*x*-axis) within the PPC cluster is not significantly correlated with behavioral logit likelihood weight (*y*-axis) (Spearman correlation: 0.156, $p = 0.475$). Each point ($N = 23$) represents one participant.

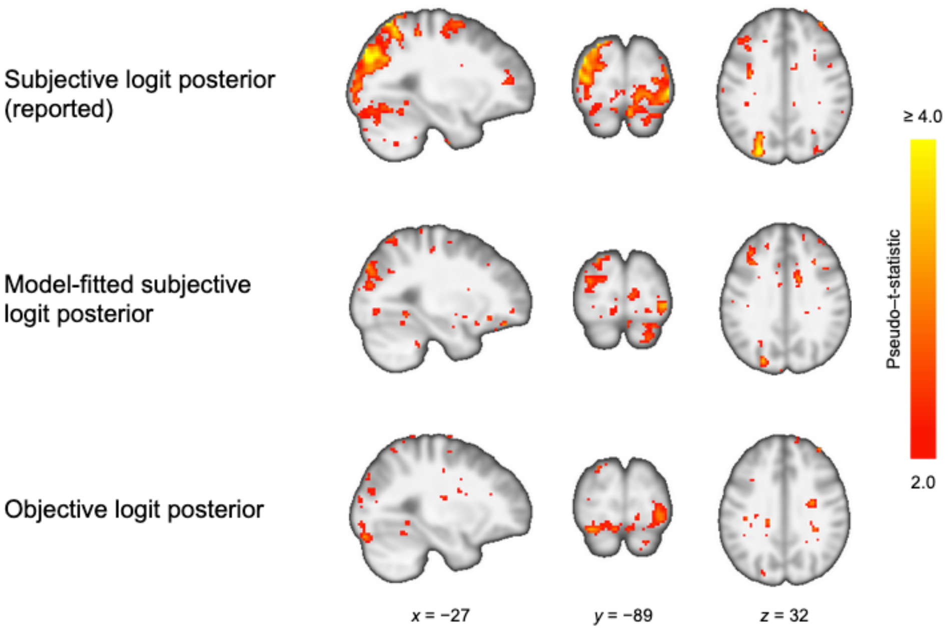


Figure S7. Activation tracking the subjective logit posterior of the questioned gallery according to the participants' reports (top row, corresponds to **Figure 3A**, WB-GLM 1); model-fitted subjective logit posterior according to **Equation 9**, the same model used to yield the regression weights in **Figure 2C** and **Figure S4A,C** (middle row, WB-GLM 4); and the objective logit posterior according to Bayes' theorem (bottom row, WB-GLM 2). All activation here is displayed at a lenient threshold for visualization purposes to allow comparison across models (uncorrected t-score ≥ 2). However, BOLD activation tracking the model-fitted subjective logit posterior and the objective logit posterior do not meet the significance threshold used for the main text.

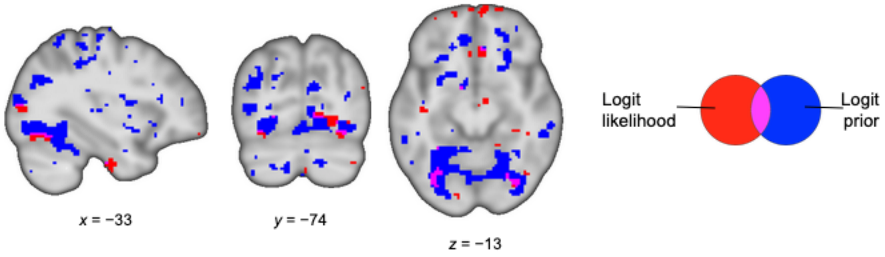


Figure S8. Whole-brain analysis indicates that clusters showing conjunction (purple) between effects of logit prior (blue) and logit likelihood (red) are sparse and small, even at a threshold (uncorrected cluster-forming height threshold: 0.05) far more lenient than the thresholds we used to define statistically significant clusters in the main text.

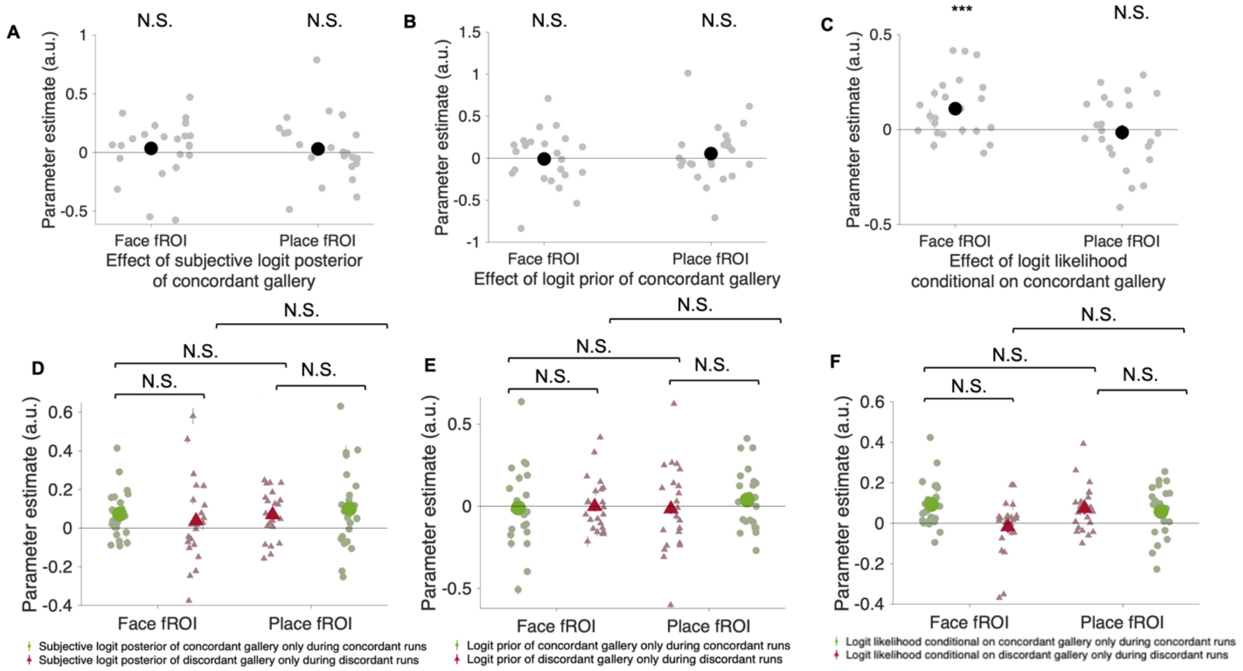


Figure S9. Face- and place-selective functional regions of interest (fROIs) show no consistent trend of probabilistic category-concordant activation.

**A–C:** In an analysis that included all trials, face- and place-selective fROIs were not significantly activated by the subjective logit posterior (**A**, duplication of **Figure 4B**) or logit prior (**B**) of their concordant galleries (i.e., the galleries corresponding to the "preferred" stimuli of that region: portrait gallery corresponding to face fROI and landscape gallery corresponding to place fROI). While the face fROI does show a positive effect of the logit likelihood conditional on the portrait gallery, the place fROI does not show a significant effect of the logit likelihood conditional on the landscape gallery (**C**). Because there were only two options (portrait gallery or landscape gallery), the probabilities of the two galleries are complementary within each analysis. Group-level statistics in black while participant-level statistics in gray. N.S.: "not significant." ***: $p < 0.001$.

**D–F:** After dividing trials by their questioned galleries (portrait or landscape), neither fROI showed preferential activation by the posterior probability of its concordant gallery and neither posterior probability had a higher parameter estimate in its concordant fMRI (**D**). The same applies to prior (**E**) and likelihood (**F**). Group-level statistics in saturated colors while participant-level statistics in pastel colors. N.S.: "not significant"

Error bars represent standard errors.

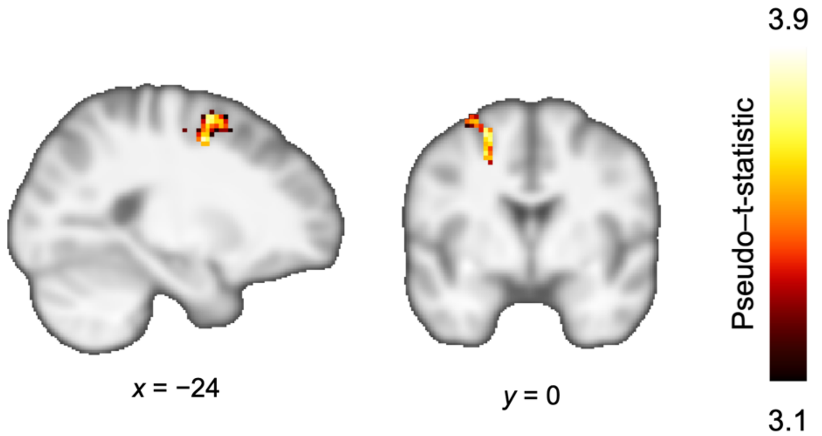

Figure S10. One cluster in the left premotor cortex shows activation by subjective logit posterior of the portrait gallery but only during portrait runs. Cluster-thresholded ($p < 0.05$, corrected for familywise error rate by permutation test) with cluster-defining height threshold of 0.001.

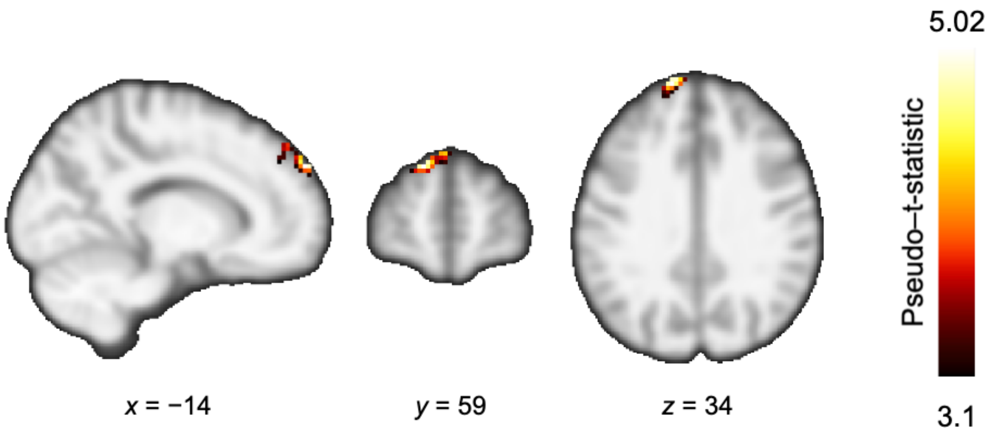


Figure S11. One cluster in the left frontal pole shows activation by logit prior of the portrait gallery during all runs. Cluster-thresholded ($p < 0.05$, corrected for familywise error rate by permutation test) with cluster-defining height threshold of 0.001.

### Supplemental Tables

Table S1. Trials in scan session. $\Pr(Q)$: Prior probability of the questioned gallery. $\theta$: Evidence strength of sample picture. $\Pr(Q|x)$: Objective posterior probability of the questioned gallery given the sample picture.

| Trial | Run | Questioned gallery | $\Pr(Q)$ | $\theta$ | $\Pr(Q\|x)$ | Penalty ($) | Hidden gallery | Sample |
|---|---|---|---|---|---|---|---|---|
| 1 | 1 | portrait | 0.53 | 0.58 | 0.45 | 20 | landscape | place |
| 2 | 1 | portrait | 0.58 | 0.80 | 0.26 | 20 | portrait | place |
| 3 | 1 | portrait | 0.50 | 0.88 | 0.88 | 20 | landscape | place |
| 4 | 1 | portrait | 0.88 | 0.91 | 0.99 | 10 | portrait | face |
| 5 | 1 | portrait | 0.49 | 0.88 | 0.12 | 10 | landscape | place |
| 6 | 1 | portrait | 0.07 | 0.90 | 0.01 | 10 | portrait | face |
| 7 | 1 | portrait | 0.49 | 0.92 | 0.92 | 10 | portrait | place |
| 8 | 1 | portrait | 0.42 | 0.80 | 0.15 | 10 | portrait | face |
| 9 | 1 | portrait | 0.48 | 0.61 | 0.59 | 20 | landscape | place |
| 10 | 1 | portrait | 0.37 | 0.88 | 0.81 | 10 | portrait | face |
| 11 | 1 | portrait | 0.61 | 0.60 | 0.70 | 20 | portrait | face |
| 12 | 1 | portrait | 0.51 | 0.88 | 0.88 | 20 | landscape | place |
| 13 | 1 | portrait | 0.50 | N/A | 0.50 | 20 | portrait | none |
| 14 | 1 | portrait | 0.39 | 0.90 | 0.07 | 20 | portrait | face |
| 15 | 1 | portrait | 0.38 | 0.81 | 0.13 | 20 | landscape | place |
| 16 | 1 | portrait | 0.47 | 0.58 | 0.55 | 10 | portrait | place |
| 17 | 1 | portrait | 0.60 | 0.59 | 0.51 | 10 | landscape | place |
| 18 | 1 | portrait | 0.87 | 0.60 | 0.91 | 10 | portrait | face |
| 19 | 1 | portrait | 0.37 | 0.91 | 0.05 | 20 | portrait | face |
| 20 | 1 | portrait | 0.51 | 0.6 | 0.61 | 10 | landscape | place |
| 21 | 1 | portrait | 0.57 | N/A | 0.57 | 10 | landscape | none |
| 22 | 1 | portrait | 0.10 | 0.60 | 0.14 | 20 | landscape | face |
| 23 | 1 | portrait | 0.92 | 0.80 | 0.98 | 20 | landscape | place |
| 24 | 1 | portrait | 0.08 | 0.80 | 0.26 | 10 | landscape | face |
| 25 | 1 | portrait | 0.12 | 0.61 | 0.08 | 20 | landscape | place |
| 26 | 1 | portrait | 0.40 | 0.77 | 0.69 | 10 | portrait | face |
| 27 | 1 | portrait | 0.60 | 0.83 | 0.24 | 20 | landscape | place |
| 28 | 1 | portrait | 0.57 | 0.88 | 0.91 | 10 | landscape | place |
| 29 | 1 | portrait | 0.40 | 0.59 | 0.32 | 20 | landscape | face |
| 30 | 1 | portrait | 0.39 | 0.87 | 0.09 | 20 | portrait | face |
| 31 | 1 | portrait | 0.48 | 0.59 | 0.57 | 10 | portrait | place |
| 32 | 1 | portrait | 0.09 | 0.88 | 0.01 | 20 | landscape | place |
| 33 | 2 | landscape | 0.51 | 0.58 | 0.59 | 10 | landscape | face |
| 34 | 2 | landscape | 0.87 | 0.92 | 0.37 | 20 | landscape | place |
| 35 | 2 | landscape | 0.61 | 0.77 | 0.32 | 20 | portrait | face |
| 36 | 2 | landscape | 0.37 | N/A | 0.37 | 10 | landscape | none |

| Trial | Run | Questioned gallery | Pr($Q$) | $\theta$ | Pr($Q\|x$) | Penalty ($) | Hidden gallery | Sample |
|---|---|---|---|---|---|---|---|---|
| 37 | 2 | landscape | 0.52 | 0.59 | 0.43 | 20 | landscape | place |
| 38 | 2 | landscape | 0.53 | 0.92 | 0.09 | 10 | landscape | place |
| 39 | 2 | landscape | 0.60 | 0.58 | 0.52 | 10 | landscape | face |
| 40 | 2 | landscape | 0.62 | 0.81 | 0.28 | 10 | landscape | face |
| 41 | 2 | landscape | 0.40 | 0.87 | 0.82 | 20 | landscape | place |
| 42 | 2 | landscape | 0.39 | 0.90 | 0.85 | 10 | landscape | place |
| 43 | 2 | landscape | 0.39 | 0.87 | 0.09 | 10 | landscape | face |
| 44 | 2 | landscape | 0.08 | 0.80 | 0.02 | 20 | landscape | place |
| 45 | 2 | landscape | 0.48 | 0.93 | 0.92 | 20 | portrait | face |
| 46 | 2 | landscape | 0.62 | 0.62 | 0.73 | 10 | landscape | place |
| 47 | 2 | landscape | 0.38 | 0.81 | 0.72 | 20 | portrait | face |
| 48 | 2 | landscape | 0.39 | 0.60 | 0.49 | 10 | landscape | place |
| 49 | 2 | landscape | 0.51 | 0.92 | 0.92 | 10 | portrait | face |
| 50 | 2 | landscape | 0.58 | 0.81 | 0.24 | 20 | landscape | place |
| 51 | 2 | landscape | 0.49 | 0.60 | 0.39 | 20 | landscape | place |
| 52 | 2 | landscape | 0.61 | 0.88 | 0.92 | 10 | portrait | face |
| 53 | 2 | landscape | 0.10 | 0.79 | 0.29 | 20 | portrait | place |
| 54 | 2 | landscape | 0.57 | 0.80 | 0.25 | 10 | portrait | face |
| 55 | 2 | landscape | 0.39 | 0.88 | 0.08 | 20 | portrait | face |
| 56 | 2 | landscape | 0.39 | N/A | 0.39 | 20 | portrait | none |
| 57 | 2 | landscape | 0.47 | 0.83 | 0.15 | 10 | portrait | place |
| 58 | 2 | landscape | 0.10 | 0.79 | 0.29 | 10 | landscape | place |
| 59 | 2 | landscape | 0.10 | 0.62 | 0.15 | 20 | landscape | place |
| 60 | 2 | landscape | 0.08 | N/A | 0.08 | 10 | portrait | none |
| 61 | 2 | landscape | 0.09 | 0.88 | 0.01 | 10 | landscape | place |
| 62 | 2 | landscape | 0.47 | 0.83 | 0.15 | 20 | landscape | place |
| 63 | 2 | landscape | 0.07 | 0.88 | 0.01 | 20 | landscape | place |
| 64 | 2 | landscape | 0.51 | 0.92 | 0.08 | 20 | portrait | place |
| 65 | 2 | landscape | 0.90 | 0.60 | 0.86 | 10 | landscape | face |
| 66 | 3 | portrait | 0.47 | 0.61 | 0.58 | 20 | landscape | face |
| 67 | 3 | portrait | 0.57 | 0.80 | 0.25 | 20 | portrait | face |
| 68 | 3 | portrait | 0.62 | 0.82 | 0.26 | 20 | landscape | place |
| 69 | 3 | portrait | 0.87 | 0.92 | 0.99 | 10 | portrait | face |
| 70 | 3 | portrait | 0.51 | 0.92 | 0.92 | 20 | portrait | face |
| 71 | 3 | portrait | 0.07 | 0.82 | 0.02 | 10 | portrait | face |
| 72 | 3 | portrait | 0.52 | 0.93 | 0.94 | 10 | landscape | place |
| 73 | 3 | portrait | 0.51 | 0.89 | 0.89 | 20 | landscape | place |
| 74 | 3 | portrait | 0.52 | 0.60 | 0.62 | 10 | landscape | place |
| 75 | 3 | portrait | 0.88 | 0.90 | 0.45 | 20 | portrait | face |

| Trial | Run | Questioned gallery | Pr($Q$) | $\theta$ | Pr($Q|x$) | Penalty ($) | Hidden gallery | Sample |
|---|---|---|---|---|---|---|---|---|
| 76 | 3 | portrait | 0.49 | 0.81 | 0.18 | 20 | portrait | face |
| 77 | 3 | portrait | 0.38 | 0.82 | 0.12 | 10 | portrait | face |
| 78 | 3 | portrait | 0.10 | 0.62 | 0.15 | 20 | portrait | place |
| 79 | 3 | portrait | 0.40 | 0.93 | 0.05 | 20 | landscape | place |
| 80 | 3 | portrait | 0.50 | 0.92 | 0.92 | 10 | landscape | place |
| 81 | 3 | portrait | 0.07 | 0.90 | 0.01 | 20 | portrait | face |
| 82 | 3 | portrait | 0.51 | 0.81 | 0.20 | 10 | landscape | place |
| 83 | 3 | portrait | 0.59 | N/A | 0.59 | 20 | portrait | none |
| 84 | 3 | portrait | 0.38 | 0.83 | 0.11 | 10 | landscape | place |
| 85 | 3 | portrait | 0.90 | 0.62 | 0.85 | 10 | portrait | place |
| 86 | 3 | portrait | 0.49 | 0.58 | 0.41 | 20 | portrait | face |
| 87 | 3 | portrait | 0.92 | N/A | 0.92 | 10 | landscape | none |
| 88 | 3 | portrait | 0.90 | 0.80 | 0.97 | 20 | landscape | place |
| 89 | 3 | portrait | 0.60 | 0.58 | 0.52 | 20 | portrait | face |
| 90 | 3 | portrait | 0.61 | 0.77 | 0.84 | 10 | landscape | place |
| 91 | 3 | portrait | 0.40 | 0.60 | 0.31 | 20 | landscape | place |
| 92 | 3 | portrait | 0.40 | 0.60 | 0.50 | 10 | landscape | face |
| 93 | 3 | portrait | 0.40 | 0.93 | 0.05 | 10 | landscape | place |
| 94 | 3 | portrait | 0.90 | 0.80 | 0.97 | 10 | landscape | place |
| 95 | 3 | portrait | 0.50 | 0.61 | 0.61 | 10 | portrait | face |
| 96 | 3 | portrait | 0.63 | 0.91 | 0.14 | 10 | landscape | place |
| 97 | 3 | portrait | 0.57 | 0.88 | 0.15 | 20 | portrait | face |
| 98 | 4 | landscape | 0.08 | 0.8 | 0.26 | 10 | portrait | face |
| 99 | 4 | landscape | 0.40 | 0.87 | 0.82 | 10 | landscape | place |
| 100 | 4 | landscape | 0.38 | 0.61 | 0.49 | 20 | landscape | place |
| 101 | 4 | landscape | 0.52 | 0.60 | 0.42 | 20 | portrait | face |
| 102 | 4 | landscape | 0.58 | 0.81 | 0.24 | 10 | landscape | place |
| 103 | 4 | landscape | 0.49 | N/A | 0.49 | 20 | portrait | none |
| 104 | 4 | landscape | 0.38 | 0.81 | 0.13 | 10 | landscape | place |
| 105 | 4 | landscape | 0.52 | 0.61 | 0.41 | 10 | landscape | place |
| 106 | 4 | landscape | 0.11 | N/A | 0.11 | 20 | portrait | none |
| 107 | 4 | landscape | 0.40 | 0.77 | 0.17 | 20 | landscape | place |
| 108 | 4 | landscape | 0.63 | 0.88 | 0.19 | 20 | landscape | face |
| 109 | 4 | landscape | 0.12 | 0.61 | 0.18 | 10 | landscape | face |
| 110 | 4 | landscape | 0.88 | 0.90 | 0.45 | 10 | landscape | face |
| 111 | 4 | landscape | 0.53 | 0.61 | 0.64 | 10 | portrait | place |
| 112 | 4 | landscape | 0.50 | 0.80 | 0.20 | 20 | landscape | place |
| 113 | 4 | landscape | 0.12 | 0.91 | 0.01 | 20 | portrait | face |
| 114 | 4 | landscape | 0.93 | 0.82 | 0.98 | 20 | portrait | face |

| Trial | Run | Questioned gallery | $\Pr(Q)$ | $\theta$ | $\Pr(Q \vert x)$ | Penalty ($) | Hidden gallery | Sample |
|---|---|---|---|---|---|---|---|---|
| 115 | 4 | landscape | 0.53 | 0.92 | 0.09 | 20 | landscape | place |
| 116 | 4 | landscape | 0.50 | 0.61 | 0.39 | 20 | landscape | place |
| 117 | 4 | landscape | 0.13 | 0.60 | 0.18 | 20 | portrait | place |
| 118 | 4 | landscape | 0.49 | 0.89 | 0.11 | 10 | landscape | place |
| 119 | 4 | landscape | 0.38 | 0.83 | 0.75 | 20 | portrait | face |
| 120 | 4 | landscape | 0.62 | 0.61 | 0.72 | 10 | portrait | face |
| 121 | 4 | landscape | 0.38 | 0.92 | 0.88 | 20 | landscape | place |
| 122 | 4 | landscape | 0.11 | N/A | 0.11 | 10 | portrait | none |
| 123 | 4 | landscape | 0.62 | 0.62 | 0.50 | 20 | landscape | face |
| 124 | 4 | landscape | 0.07 | 0.88 | 0.36 | 10 | landscape | face |
| 125 | 4 | landscape | 0.62 | 0.92 | 0.95 | 10 | portrait | face |
| 126 | 4 | landscape | 0.40 | 0.83 | 0.76 | 10 | landscape | place |
| 127 | 4 | landscape | 0.50 | 0.80 | 0.20 | 10 | landscape | place |
| 128 | 4 | landscape | 0.90 | 0.62 | 0.94 | 10 | portrait | face |
| 129 | 4 | landscape | 0.50 | 0.92 | 0.08 | 20 | portrait | face |
| 130 | 4 | landscape | 0.50 | 0.88 | 0.88 | 10 | portrait | place |

Table S2. Mean and standard deviation of regressors in the models of subjective logit posterior in **Figure S3**.

| Regressor | | Model(s) | Mean | Sample standard deviation |
|---|---|---|---|---|
| Name | Symbol | | | |
| Objective logit posterior | $\log\left(\frac{\Pr(Q \vert x)}{1-\Pr(Q \vert x)}\right)$ | Base | 0.0221 | 2.0944 |
| Logit prior | $\log\left(\frac{\Pr(Q)}{1-\Pr(Q)}\right)$ | Reduced, extended | 0.0063 | 1.1939 |
| Logit likelihood | $\log\left(\frac{\Pr(x \vert Q)}{1-\Pr(x \vert Q)}\right)$ | Reduced, extended | 0.0157 | 1.6012 |
| Inaccuracy penalty | $W$ | Extended | $15.00 | $5.0009 |
| Initial slider position | $H$ | Extended | 981.5891 | 551.0087 |

Table S3. Fixed-effects regression weights in the model of subjective logit posterior as a function of objective logit posterior (**Equation 7**). Regressors were z-scored before the model was fit. *DF*: Degree of freedom. *p*: *p*-value.

| Regressor | Regressor weight | | Standard error | | $T$-statistic | DF | $p$ | |
|---|---|---|---|---|---|---|---|---|
| | Z | M | Z | M | | | Z | M |
| Intercept | 0.072 | 0.072 | 0.028 | 0.028 | 2.571 | 2,748 | 0.010 | 0.001 |
| Objective logit posterior | 1.171 | 0.560 | 0.068 | 0.033 | 17.170 | 2,748 | < 0.001 | < 0.001 |

Table S4. Fixed-effects regression weights in the model of subjective logit posterior as a function of logit prior and logit likelihood (**Equation 8**). Z: statistics from regression with z-scored predictors; corresponds to **Figure**

**S4D**. M: statistics from regression with mean-centered (but not z-scored) predictors; corresponds to **Figure S4B**.
*DF*: Degree of freedom. *p*: *p*-value.

| Regressor | Regressor weight | | Standard error | | *T*-statistic | DF | *p* | |
|---|---|---|---|---|---|---|---|---|
| | Z | M | Z | M | | | Z | M |
| Intercept | 0.073 | 0.073 | 0.028 | 0.028 | 2.570 | 2,747 | 0.010 | 0.010 |
| Logit prior | 0.780 | 0.654 | 0.056 | 0.047 | 13.986 | 2,747 | < 0.001 | < 0.001 |
| Logit likelihood | 0.806 | 0.503 | 0.079 | 0.049 | 10.260 | 2,747 | < 0.001 | < 0.001 |

Table S5. Fixed-effects regression weights in the model of subjective logit posterior as a function of logit prior, logit likelihood, and nuisance regressors (**Equation 9**). Regressors were z-scored before the model was fit; corresponds to **Figure S4C** and **Figure 2C**. *DF*: Degree of freedom. *p*: -value.

| Regressor | Regression weight | | Standard error | | *T*-statistic | DF | *p* | |
|---|---|---|---|---|---|---|---|---|
| | Z | M | Z | M | | | Z | M |
| Intercept | 0.073 | 0.073 | 0.028 | 0.028 | 2.583 | 2,745 | 0.010 | 0.010 |
| Logit prior | 0.789 | 0.661 | 0.056 | 0.047 | 14.181 | 2,745 | < 0.001 | < 0.001 |
| Logit likelihood | 0.804 | 0.502 | 0.078 | 0.049 | 10.333 | 2,745 | < 0.001 | < 0.001 |
| Inaccuracy penalty | 0.040 | 0.008 | 0.012 | 0.002 | 3.205 | 2,745 | 0.001 | 0.001 |
| Initial slider position | 0.035 | $6.4\times10^{-5}$ | 0.013 | $2.4\times10^{-5}$ | 2.658 | 2,745 | 0.008 | 0.008 |

Table S6. Peak activations in the cluster (467 voxels) in left posterior parietal cortex (PPC) that was significantly activated by subjective logit posterior (**Figure 3A**). Anatomical regions according to the JuBrain/SPM Anatomy Toolbox 3.0[59–61] and the Oxford-Harvard Atlas[62].

| Pseudo–*t*-statistic | MNI Coordinates | | | Anatomical region | |
|---|---|---|---|---|---|
| | *x* | *y* | *z* | JuBrain | Oxford/Harvard Atlas |
| 5.07 | −19 | −67 | 49 | 7A (SPL) | Lateral Occipital Cortex, superior division |
| 4.70 | −17 | −58 | 73 | N/A | Superior Parietal Lobule |
| 4.53 | −24 | −65 | 66 | 7A (SPL) | Lateral Occipital Cortex, superior division |
| 4.50 | −27 | −84 | 32 | hIP4 (IPS) | Lateral Occipital Cortex, superior division |
| 4.12 | −36 | −89 | 25 | hOc4lp | Lateral Occipital Cortex, superior division |
| 4.12 | −34 | −65 | 61 | 7A (SPL) | Lateral Occipital Cortex, superior division |
| 4.03 | −24 | −60 | 59 | 7A (SPL) | Lateral Occipital Cortex, superior division |
| 3.98 | −34 | −94 | 18 | hOc4lp | Occipital Pole |
| 3.97 | −27 | −77 | 35 | hIP5 (IPS) | Lateral Occipital Cortex, superior division |
| 3.94 | −36 | −94 | 6 | hOc4lp | Occipital Pole |

Table S7. Parameter estimates of activation of PPC cluster (**Figure 3A**) for the quadratic and linear terms for subjective logit posterior along with nuisance regressors. *DF*: Degrees of freedom. *p*: *p*-value.

| Regressor | Parameter estimate | Standard error | T-statistic | DF | *p* |
|---|---|---|---|---|---|
| $(\text{Subjective logit posterior})^2$ | 0.077 | 0.042 | 1.81 | 42,960 | 0.070 |
| Subjective logit posterior | 0.264 | 0.057 | 4.616 | 42,960 | < 0.001 |
| Inaccuracy penalty | −0.012 | 0.011 | −1.110 | 42,960 | 0.267 |

| | | | | | |
|---|---|---|---|---|---|
| Slider displacement | 0.000 | 0.005 | −0.003 | 42,960 | 0.997 |

Table S8. Parameter estimates of activation of PPC cluster (**Figure 3A**) for logit prior, logit likelihood, and nuisance regressors. Corresponds to statistics in **Figure 3C**. *DF*: Degrees of freedom. *p*: *p*-value.

| Regressor | Parameter estimate | Standard error | T-statistic | DF | *p* |
|---|---|---|---|---|---|
| Logit prior | 0.131 | 0.063 | 2.100 | 42,960 | 0.039 |
| Logit likelihood | 0.202 | 0.039 | 3.139 | 42,960 | < 0.001 |
| Inaccuracy penalty | −0.011 | 0.012 | −1.580 | 42,960 | 0.341 |
| Slider displacement | 0.000 | 0.004 | 0.027 | 42,960 | 0.988 |

Table S9. Parameter estimates of activation of PPC cluster (**Figure 3A**) for logit prior, subjective logit posterior, and nuisance regressors. Corresponds to statistics in **Figure S5A**. *DF*: Degrees of freedom. *p*: *p*-value.

| Regressor | Parameter estimate | Standard error | *T*-statistic | DF | *p* |
|---|---|---|---|---|---|
| Logit prior | −0.084 | 0.079 | −1.060 | 42,960 | 0.289 |
| Subjective logit posterior | 0.386 | 0.072 | 5.356 | 42,960 | < 0.001 |
| Inaccuracy penalty | −0.013 | 0.012 | −1.123 | 42,960 | 0.262 |
| Slider displacement | 0.000 | 0.006 | −0.004 | 42,960 | 0.996 |

Table S10. Parameter estimates of activation of PPC cluster (**Figure 3A**) for logit likelihood, subjective logit posterior, and nuisance regressors. Corresponds to **Figure S5B**. *DF*: Degrees of freedom. *p*: *p*-value.

| Regressor | Parameter estimate | Standard error | *T*-statistic | DF | *p* |
|---|---|---|---|---|---|
| Logit likelihood | 0.036 | 0.073 | 0.492 | 42,960 | 0.623 |
| Subjective logit posterior | 0.331 | 0.080 | 4.136 | 42,960 | < 0.001 |
| Inaccuracy penalty | −0.014 | 0.011 | −1.328 | 42,960 | 0.184 |
| Slider displacement | 0.000 | 0.006 | 0.000 | 42,960 | 0.999 |

Table S11. Fixed-effects parameter estimates for activation of face and place functional regions of interest (fROIs) by the subjective logit posterior of the gallery signaled by their preferred stimuli. Corresponds to **Figure 4B** and **Figure S9A**. *DF*: Degrees of freedom. *p*: *p*-value.

| fROI | Regressor | Parameter estimate | Standard error | *T*-statistic | DF | *p* |
|---|---|---|---|---|---|---|
| Face-selective | Subjective logit posterior of portrait gallery across all runs | 0.035 | 0.052 | 0.671 | 569,318 | 0.502 |
| | Inaccuracy penalty | −0.003 | 0.006 | −0.546 | 569,318 | 0.585 |
| | Slider displacement | 0.000 | 0.003 | 0.016 | 569,318 | 0.987 |

| | | | | | | |
|---|---|---|---|---|---|---|
| Place-selective | Subjective logit posterior of landscape gallery across all runs | 0.030 | 0.055 | 0.545 | 569,318 | 0.586 |
| | Inaccuracy penalty | −0.001 | 0.005 | −0.188 | 569,318 | 0.851 |
| | Slider displacement | 0.000 | 0.002 | −0.016 | 569,318 | 0.987 |

Table S12. Fixed-effects parameter estimates for activation of face and place functional regions of interest (fROIs) by the logit prior of the gallery signaled by their preferred stimuli and logit likelihood conditional on the gallery signaled by their preferred stimuli. Corresponds to **Figure S9B–C**. *DF*: Degrees of freedom. *p*: *p*-value.

| fROI | Regressor | Parameter estimate | Standard error | *T*-statistic | DF | *p* |
|---|---|---|---|---|---|---|
| Face-selective | Logit prior of portrait gallery across all runs | −0.009 | 0.068 | −0.130 | 853,978 | 0.896 |
| | Logit likelihood conditional on portrait gallery across all runs | 0.110 | 0.033 | 3.400 | 853,978 | < 0.001 |
| | Inaccuracy penalty | −0.004 | 0.006 | −0.587 | 853,978 | 0.557 |
| | Slider displacement | $7\times10^{-5}$ | 0.003 | 0.025 | 853,978 | 0.980 |
| Place-selective | Logit prior of landscape gallery across all runs | 0.055 | 0.071 | 0.777 | 853,978 | 0.437 |
| | Logit likelihood conditional on landscape gallery across all runs | −0.015 | 0.038 | −0.384 | 853,978 | 0.701 |
| | Inaccuracy penalty | −0.001 | 0.005 | −0.233 | 853,978 | 0.816 |
| | Slider displacement | $-7\times10^{-6}$ | 0.002 | $-6\times10^{-4}$ | 853,978 | > 0.999 |

Table S13. Three-way ANOVA for effects of participant, contrast, and functional region of interest (fROI) on mean signal of the face and place fROIs, with focus on the subjective logit posterior.
Predictors:
"participant": identifier for each participant (23 total)
"contrast": (1) subjective logit posterior of portrait gallery only during portrait runs, (2) subjective logit posterior of landscape gallery only during landscape runs, (3) inaccuracy penalty, (4) slider displacement
"fROI": (1) face-selective fROI and (2) place-selective fROI
Pairwise comparisons in **Table S14**.

| Predictor | Sum of Squares | DF | Mean Squares | *F*-statistic | *p* |
|---|---|---|---|---|---|
| participant | 0.571 | 22 | 0.026 | 1.855 | 0.016 |
| contrast | 0.226 | 3 | 0.075 | 5.391 | 0.001 |
| fROI | 0.010 | 1 | 0.010 | 0.686 | 0.409 |
| contrast*fROI | 0.034 | 3 | 0.011 | 0.811 | 0.490 |
| Error | 2.156 | 154 | 0.014 | | |
| Total | 2.997 | 183 | | | |

Table S14. Pairwise comparisons for predictors of contrast and functional region of interest (fROI) from the ANOVA in **Table S13** (Tukey Honest Significant Difference Test). Rows in black text represent comparisons plotted in **Figure 4C** and **Figure S9D**. Rows in gray text represent unplotted comparisons. All contrasts were positive.

| Predictor 1 | | Predictor 2 | | Lower CI | Difference | Upper CI | *p* |
|---|---|---|---|---|---|---|---|
| Contrast | fROI | Contrast | fROI | | | | |
| Logit posterior of portrait gallery only on portrait runs | Face | Logit posterior of landscape gallery only on landscape runs | Face | −0.071 | 0.034 | 0.140 | 0.976 |
| Logit posterior of portrait gallery only on portrait runs | Face | Penalty | Face | −0.033 | 0.072 | 0.178 | 0.431 |
| Logit posterior of portrait gallery only on portrait runs | Face | Cursor displacement | Face | −0.033 | 0.073 | 0.178 | 0.427 |
| Logit posterior of portrait gallery only on portrait runs | Face | Logit posterior of portrait gallery only on portrait runs | Place | −0.103 | 0.003 | 0.109 | 1.000 |
| Logit posterior of portrait gallery only on portrait runs | Face | Logit posterior of landscape gallery only on landscape runs | Place | −0.133 | −0.027 | 0.079 | 0.994 |
| Logit posterior of portrait gallery only on portrait runs | Face | Penalty | Place | −0.033 | 0.073 | 0.179 | 0.417 |
| Logit posterior of portrait gallery only on portrait runs | Face | Cursor displacement | Place | −0.033 | 0.073 | 0.178 | 0.426 |
| Logit posterior of landscape gallery only on landscape runs | Face | Penalty | Face | −0.068 | 0.038 | 0.144 | 0.959 |
| Logit posterior of landscape gallery only on landscape runs | Face | Cursor displacement | Face | −0.068 | 0.038 | 0.144 | 0.958 |
| Logit posterior of landscape gallery only on landscape runs | Face | Logit posterior of portrait gallery only on portrait runs | Place | −0.137 | −0.031 | 0.074 | 0.986 |
| Logit posterior of landscape gallery only on landscape runs | Face | Logit posterior of landscape gallery only on landscape runs | Place | −0.167 | −0.062 | 0.044 | 0.645 |
| Logit posterior of landscape gallery only on landscape runs | Face | Penalty | Place | −0.067 | 0.039 | 0.144 | 0.955 |
| Logit posterior of landscape gallery only on landscape runs | Face | Cursor displacement | Place | −0.068 | 0.038 | 0.144 | 0.958 |
| Penalty | Face | Cursor displacement | Face | −0.106 | 0.000 | 0.106 | 1.000 |
| Penalty | Face | Logit posterior of portrait gallery only on portrait runs | Place | −0.175 | −0.069 | 0.036 | 0.488 |
| Penalty | Face | Logit posterior of landscape gallery only on landscape runs | Place | −0.205 | −0.100 | 0.006 | 0.083 |
| Penalty | Face | Penalty | Place | −0.105 | 0.001 | 0.107 | 1.000 |
| Penalty | Face | Cursor displacement | Place | −0.105 | 0.000 | 0.106 | 1.000 |
| Cursor displacement | Face | Logit posterior of portrait gallery only on portrait runs | Place | −0.175 | −0.070 | 0.036 | 0.484 |
| Cursor displacement | Face | Logit posterior of landscape gallery only on landscape runs | Place | −0.205 | −0.100 | 0.006 | 0.081 |
| Cursor displacement | Face | Penalty | Place | −0.105 | 0.001 | 0.106 | 1.000 |

| Predictor 1 | | Predictor 2 | | Lower CI | Difference | Upper CI | $p$ |
|---|---|---|---|---|---|---|---|
| Contrast | fROI | Contrast | fROI | | | | |
| Cursor displacement | Face | Cursor displacement | Place | −0.106 | 0.000 | 0.106 | 1.000 |
| Logit posterior of portrait gallery only on portrait runs | Place | Logit posterior of landscape gallery only on landscape runs | Place | −0.136 | −0.030 | 0.076 | 0.989 |
| Logit posterior of portrait gallery only on portrait runs | Place | Penalty | Place | −0.036 | 0.070 | 0.176 | 0.473 |
| Logit posterior of portrait gallery only on portrait runs | Place | Cursor displacement | Place | −0.036 | 0.070 | 0.175 | 0.483 |
| Logit posterior of landscape gallery only on landscape runs | Place | Penalty | Place | −0.005 | 0.100 | 0.206 | 0.078 |
| Logit posterior of landscape gallery only on landscape runs | Place | Cursor displacement | Place | −0.006 | 0.100 | 0.206 | 0.081 |
| Penalty | Place | Cursor displacement | Place | −0.106 | −0.001 | 0.105 | 1.000 |

Table S15. Three-way ANOVA for effects of participant, contrast, and functional region of interest (fROI) on mean signal of the face and place fROIs, with focus on logit prior and logit likelihood.
Predictors:
"participant": identifier for each participant (23 total)
"contrast": (1) logit prior of portrait gallery only during portrait runs, (2) logit prior of landscape gallery only during landscape runs, (3) logit likelihood conditional on portrait gallery only during portrait runs, (4) logit likelihood conditional on landscape gallery only during landscape gallery, (5) inaccuracy penalty, (6) slider displacement
"fROI": (1) face-selective fROI and (2) place-selective fROI
Pairwise comparisons in **Table S16**.

| Predictor | Sum of Squares | DF | Mean Squares | $F$-statistic | $p$ |
|---|---|---|---|---|---|
| participant | 0.731 | 22 | 0.033 | 1.702 | 0.029 |
| contrast | 0.272 | 5 | 0.054 | 2.784 | 0.018 |
| fROI | 0.016 | 1 | 0.016 | 0.797 | 0.373 |
| contrast*fROI | 0.074 | 5 | 0.015 | 0.758 | 0.581 |
| Error | 4.725 | 242 | 0.020 | | |
| Total | 5.817 | 275 | | | |

Table S16. Pairwise comparisons for predictors of contrast and functional region of interest (fROI) from the ANOVA for effects of logit prior and logit likelihood in **Table S15** (Tukey Honest Significant Difference Test). Rows in black text represent comparisons plotted in **Figure S9E–F**. Rows in gray text represent unplotted comparisons. All contrasts were positive.

| Predictor 1 | | Predictor 2 | | Lower CI | Difference | Upper CI | $p$ |
|---|---|---|---|---|---|---|---|
| Contrast | fROI | Contrast | fROI | | | | |
| Logit prior of portrait gallery only on portrait runs | Face | Logit prior of landscape gallery only on landscape runs | Face | −0.144 | −0.009 | 0.126 | 1.000 |
| Logit prior of portrait gallery only on portrait runs | Face | Logit likelihood conditional on portrait gallery only on portrait runs | Face | −0.238 | −0.103 | 0.031 | 0.336 |
| Logit prior of portrait gallery only on portrait runs | Face | Logit likelihood conditional on landscape gallery only on landscape runs | Face | −0.128 | 0.007 | 0.142 | 1.000 |

| Predictor 1 | | Predictor 2 | | Lower CI | Difference | Upper CI | $p$ |
|---|---|---|---|---|---|---|---|
| Contrast | fROI | Contrast | fROI | | | | |
| Logit prior of portrait gallery only on portrait runs | Face | Logit prior of landscape gallery only on landscape runs | Face | −0.144 | −0.009 | 0.126 | 1.000 |
| Logit prior of portrait gallery only on portrait runs | Face | Logit likelihood conditional on portrait gallery only on portrait runs | Face | −0.238 | −0.103 | 0.031 | 0.336 |
| Logit prior of portrait gallery only on portrait runs | Face | Penalty | Face | −0.144 | −0.009 | 0.126 | 1.000 |
| Logit prior of portrait gallery only on portrait runs | Face | Slider displacement | Face | −0.145 | −0.010 | 0.124 | 1.000 |
| Logit prior of portrait gallery only on portrait runs | Face | Logit prior of portrait gallery only on portrait runs | Place | −0.128 | 0.006 | 0.141 | 1.000 |
| Logit prior of portrait gallery only on portrait runs | Face | Logit prior of landscape gallery only on landscape runs | Place | −0.184 | −0.049 | 0.086 | 0.990 |
| Logit prior of portrait gallery only on portrait runs | Face | Logit likelihood conditional on portrait gallery only on portrait runs | Place | −0.218 | −0.083 | 0.051 | 0.678 |
| Logit prior of portrait gallery only on portrait runs | Face | Logit likelihood conditional on landscape gallery only on landscape runs | Place | −0.203 | −0.069 | 0.066 | 0.882 |
| Logit prior of portrait gallery only on portrait runs | Face | Penalty | Place | −0.144 | −0.009 | 0.125 | 1.000 |
| Logit prior of portrait gallery only on portrait runs | Face | Slider displacement | Place | −0.145 | −0.010 | 0.124 | 1.000 |
| Logit prior of landscape gallery only on landscape runs | Face | Logit likelihood conditional on portrait gallery only on portrait runs | Face | −0.229 | −0.094 | 0.040 | 0.484 |
| Logit prior of landscape gallery only on landscape runs | Face | Logit likelihood conditional on landscape gallery only on landscape runs | Face | −0.119 | 0.016 | 0.151 | 1.000 |
| Logit prior of landscape gallery only on landscape runs | Face | Penalty | Face | −0.135 | 0.000 | 0.135 | 1.000 |
| Logit prior of landscape gallery only on landscape runs | Face | Slider displacement | Face | −0.136 | −0.001 | 0.133 | 1.000 |
| Logit prior of landscape gallery only on landscape runs | Face | Logit prior of portrait gallery only on portrait runs | Place | −0.119 | 0.015 | 0.150 | 1.000 |
| Logit prior of landscape gallery only on landscape runs | Face | Logit prior of landscape gallery only on landscape runs | Place | −0.175 | −0.040 | 0.095 | 0.998 |
| Logit prior of landscape gallery only on landscape runs | Face | Logit likelihood conditional on portrait gallery only on portrait runs | Place | −0.209 | −0.074 | 0.060 | 0.815 |
| Logit prior of landscape gallery only on landscape runs | Face | Logit likelihood conditional on landscape gallery only on landscape runs | Place | −0.195 | −0.060 | 0.075 | 0.953 |

| Predictor 1 | | Predictor 2 | | Lower CI | Difference | Upper CI | $p$ |
|---|---|---|---|---|---|---|---|
| Contrast | fROI | Contrast | fROI | | | | |
| Logit prior of portrait gallery only on portrait runs | Face | Logit prior of landscape gallery only on landscape runs | Face | −0.144 | −0.009 | 0.126 | 1.000 |
| Logit prior of portrait gallery only on portrait runs | Face | Logit likelihood conditional on portrait gallery only on portrait runs | Face | −0.238 | −0.103 | 0.031 | 0.336 |
| Logit prior of landscape gallery only on landscape runs | Face | Penalty | Place | −0.135 | 0.000 | 0.134 | 1.000 |
| Logit prior of landscape gallery only on landscape runs | Face | Slider displacement | Place | −0.136 | −0.001 | 0.133 | 1.000 |
| Logit likelihood conditional on portrait gallery only on portrait runs | Face | Logit likelihood conditional on landscape gallery only on landscape runs | Face | −0.024 | 0.110 | 0.245 | 0.238 |
| Logit likelihood conditional on portrait gallery only on portrait runs | Face | Penalty | Face | −0.040 | 0.094 | 0.229 | 0.484 |
| Logit likelihood conditional on portrait gallery only on portrait runs | Face | Slider displacement | Face | −0.042 | 0.093 | 0.228 | 0.510 |
| Logit likelihood conditional on portrait gallery only on portrait runs | Face | Logit prior of portrait gallery only on portrait runs | Place | −0.025 | 0.109 | 0.244 | 0.248 |
| Logit likelihood conditional on portrait gallery only on portrait runs | Face | Logit prior of landscape gallery only on landscape runs | Place | −0.080 | 0.054 | 0.189 | 0.977 |
| Logit likelihood conditional on portrait gallery only on portrait runs | Face | Logit likelihood conditional on portrait gallery only on portrait runs | Place | −0.115 | 0.020 | 0.155 | 1.000 |
| Logit likelihood conditional on portrait gallery only on portrait runs | Face | Logit likelihood conditional on landscape gallery only on landscape runs | Place | −0.100 | 0.034 | 0.169 | 1.000 |
| Logit likelihood conditional on portrait gallery only on portrait runs | Face | Penalty | Place | −0.041 | 0.094 | 0.229 | 0.492 |
| Logit likelihood conditional on portrait gallery only on portrait runs | Face | Slider displacement | Place | −0.042 | 0.093 | 0.228 | 0.510 |
| Logit likelihood conditional on landscape gallery only on landscape runs | Face | Penalty | Face | −0.151 | −0.016 | 0.119 | 1.000 |
| Logit likelihood conditional on landscape gallery only on landscape runs | Face | Slider displacement | Face | −0.152 | −0.017 | 0.117 | 1.000 |
| Logit likelihood conditional on landscape gallery only on landscape runs | Face | Logit prior of portrait gallery only on portrait runs | Place | −0.135 | −0.001 | 0.134 | 1.000 |
| Logit likelihood conditional on landscape gallery only on landscape runs | Face | Logit prior of landscape gallery only on landscape runs | Place | −0.191 | −0.056 | 0.079 | 0.971 |

| Predictor 1 | | Predictor 2 | | Lower CI | Difference | Upper CI | $p$ |
|---|---|---|---|---|---|---|---|
| Contrast | fROI | Contrast | fROI | | | | |
| Logit prior of portrait gallery only on portrait runs | Face | Logit prior of landscape gallery only on landscape runs | Face | −0.144 | −0.009 | 0.126 | 1.000 |
| Logit prior of portrait gallery only on portrait runs | Face | Logit likelihood conditional on portrait gallery only on portrait runs | Face | −0.238 | −0.103 | 0.031 | 0.336 |
| Logit likelihood conditional on landscape gallery only on landscape runs | Face | Logit likelihood conditional on portrait gallery only on portrait runs | Place | −0.225 | −0.090 | 0.044 | 0.554 |
| Logit likelihood conditional on landscape gallery only on landscape runs | Face | Logit likelihood conditional on landscape gallery only on landscape runs | Place | −0.211 | −0.076 | 0.059 | 0.795 |
| Logit likelihood conditional on landscape gallery only on landscape runs | Face | Penalty | Place | −0.151 | −0.016 | 0.118 | 1.000 |
| Logit likelihood conditional on landscape gallery only on landscape runs | Face | Slider displacement | Place | −0.152 | −0.017 | 0.117 | 1.000 |
| Penalty | Face | Slider displacement | Face | −0.136 | −0.001 | 0.133 | 1.000 |
| Penalty | Face | Logit prior of portrait gallery only on portrait runs | Place | −0.119 | 0.015 | 0.150 | 1.000 |
| Penalty | Face | Logit prior of landscape gallery only on landscape runs | Place | −0.175 | −0.040 | 0.095 | 0.998 |
| Penalty | Face | Logit likelihood conditional on portrait gallery only on portrait runs | Place | −0.209 | −0.074 | 0.060 | 0.815 |
| Penalty | Face | Logit likelihood conditional on landscape gallery only on landscape runs | Place | −0.195 | −0.060 | 0.075 | 0.953 |
| Penalty | Face | Penalty | Place | −0.135 | 0.000 | 0.134 | 1.000 |
| Penalty | Face | Slider displacement | Place | −0.136 | −0.001 | 0.133 | 1.000 |
| Slider displacement | Face | Logit prior of portrait gallery only on portrait runs | Place | −0.118 | 0.017 | 0.151 | 1.000 |
| Slider displacement | Face | Logit prior of landscape gallery only on landscape runs | Place | −0.173 | −0.039 | 0.096 | 0.999 |
| Slider displacement | Face | Logit likelihood conditional on portrait gallery only on portrait runs | Place | −0.208 | −0.073 | 0.062 | 0.834 |
| Slider displacement | Face | Logit likelihood conditional on landscape gallery only on landscape runs | Place | −0.193 | −0.058 | 0.076 | 0.960 |
| Slider displacement | Face | Penalty | Place | −0.134 | 0.001 | 0.136 | 1.000 |
| Slider displacement | Face | Slider displacement | Place | −0.135 | 0.000 | 0.135 | 1.000 |
| Logit prior of portrait gallery only on portrait runs | Place | Logit prior of landscape gallery only on landscape runs | Place | −0.190 | −0.055 | 0.079 | 0.974 |

| Predictor 1 | | Predictor 2 | | Lower CI | Difference | Upper CI | *p* |
|---|---|---|---|---|---|---|---|
| Contrast | fROI | Contrast | fROI | | | | |
| Logit prior of portrait gallery only on portrait runs | Face | Logit prior of landscape gallery only on landscape runs | Face | −0.144 | −0.009 | 0.126 | 1.000 |
| Logit prior of portrait gallery only on portrait runs | Face | Logit likelihood conditional on portrait gallery only on portrait runs | Face | −0.238 | −0.103 | 0.031 | 0.336 |
| Logit prior of portrait gallery only on portrait runs | Place | Logit likelihood conditional on portrait gallery only on portrait runs | Place | −0.224 | −0.090 | 0.045 | 0.568 |
| Logit prior of portrait gallery only on portrait runs | Place | Logit likelihood conditional on landscape gallery only on landscape runs | Place | −0.210 | −0.075 | 0.060 | 0.807 |
| Logit prior of portrait gallery only on portrait runs | Place | Penalty | Place | −0.150 | −0.016 | 0.119 | 1.000 |
| Logit prior of portrait gallery only on portrait runs | Place | Slider displacement | Place | −0.151 | −0.017 | 0.118 | 1.000 |
| Logit prior of landscape gallery only on landscape runs | Place | Logit likelihood conditional on portrait gallery only on portrait runs | Place | −0.169 | −0.034 | 0.100 | 1.000 |
| Logit prior of landscape gallery only on landscape runs | Place | Logit likelihood conditional on landscape gallery only on landscape runs | Place | −0.155 | −0.020 | 0.115 | 1.000 |
| Logit prior of landscape gallery only on landscape runs | Place | Penalty | Place | −0.095 | 0.040 | 0.174 | 0.998 |
| Logit prior of landscape gallery only on landscape runs | Place | Slider displacement | Place | −0.096 | 0.039 | 0.173 | 0.999 |
| Logit likelihood conditional on portrait gallery only on portrait runs | Place | Logit likelihood conditional on landscape gallery only on landscape runs | Place | −0.120 | 0.015 | 0.149 | 1.000 |
| Logit likelihood conditional on portrait gallery only on portrait runs | Place | Penalty | Place | −0.061 | 0.074 | 0.209 | 0.821 |
| Logit likelihood conditional on portrait gallery only on portrait runs | Place | Slider displacement | Place | −0.062 | 0.073 | 0.208 | 0.833 |
| Logit likelihood conditional on landscape gallery only on landscape runs | Place | Penalty | Place | −0.075 | 0.059 | 0.194 | 0.955 |
| Logit likelihood conditional on landscape gallery only on landscape runs | Place | Slider displacement | Place | −0.076 | 0.058 | 0.193 | 0.960 |
| Penalty | Place | Slider displacement | Place | −0.136 | −0.001 | 0.134 | 1.000 |

Table S17. Activation by subjective logit posterior by gallery category.

| Contrast | Significant Cluster(s)? |
|---|---|
| Subjective logit posterior of portrait gallery | **Figure S10** |
| Subjective logit posterior of landscape gallery | No |

Table S18. Activation by logit prior and logit likelihood by gallery category.

| Contrast | Significant Cluster(s)? |
| --- | --- |
| Logit prior of portrait gallery | **Figure S11** |
| Logit prior of landscape gallery | No |
| Logit likelihood conditional on portrait gallery | No |
| Logit likelihood conditional on landscape gallery | No |

### Details on fMRI Preprocessing

Results included in this manuscript come from preprocessing performed using *fMRIPrep* 1.5.0rc1 (Esteban, Markiewicz, et al. (2018); Esteban, Blair, et al. (2018); RRID:SCR_016216), which is based on *Nipype* 1.2.0 (Gorgolewski et al. (2011); Gorgolewski et al. (2018); RRID:SCR_002502).

#### Anatomical data preprocessing

The T1-weighted (T1w) image was corrected for intensity non-uniformity (INU) with N4BiasFieldCorrection (Tustison et al. 2010), distributed with ANTs 2.2.0 (Avants et al. 2008, RRID:SCR_004757), and used as T1w-reference throughout the workflow. The T1w-reference was then skull-stripped with a *Nipype* implementation of the antsBrainExtraction.sh workflow (from ANTs), using OASIS30ANTs as target template. Brain tissue segmentation of cerebrospinal fluid (CSF), white-matter (WM) and gray-matter (GM) was performed on the brain-extracted T1w using fast (FSL 5.0.9, RRID:SCR_002823, Zhang, Brady, and Smith 2001). Brain surfaces were reconstructed using recon-all (FreeSurfer 6.0.1, RRID:SCR_001847, Dale, Fischl, and Sereno 1999), and the brain mask estimated previously was refined with a custom variation of the method to reconcile ANTs-derived and FreeSurfer-derived segmentations of the cortical gray-matter of Mindboggle (RRID:SCR_002438, Klein et al. 2017). Volume-based spatial normalization to one standard space (MNI152NLin2009cAsym) was performed through nonlinear registration with antsRegistration (ANTs 2.2.0), using brain-extracted versions of both T1w reference and the T1w template. The following template was selected for spatial normalization: *ICBM 152 Nonlinear Asymmetrical template version 2009c* [Fonov et al. (2009), RRID:SCR_008796; TemplateFlow ID: MNI152NLin2009cAsym].

Functional data preprocessing

For each of the 10 BOLD runs found per subject (across all tasks and sessions), the following preprocessing was performed. First, a reference volume and its skull-stripped version were generated using a custom methodology of *fMRIPrep*. A deformation field to correct for susceptibility distortions was estimated based on two echo-planar imaging (EPI) references with opposing phase-encoding directions, using 3dQwarp Cox and Hyde (1997) (AFNI 20160207). Based on the estimated susceptibility distortion, an unwarped BOLD reference was calculated for a more accurate co-registration with the anatomical reference. The BOLD reference was then co-registered to the T1w reference using bbregister (FreeSurfer) which implements boundary-based registration (Greve and Fischl 2009). Co-registration was configured with six degrees of freedom. Head-motion parameters with respect to the BOLD reference (transformation matrices, and six corresponding rotation and translation parameters) are estimated before any spatiotemporal filtering using mcflirt (FSL 5.0.9, Jenkinson et al. 2002). BOLD runs were slice-time corrected using 3dTshift from AFNI 20160207 (Cox and Hyde 1997, RRID:SCR_005927). The BOLD time-series, were resampled to surfaces on the following spaces: *fsaverage5*. The BOLD time-series (including slice-timing correction when applied) were resampled onto their original, native space by applying a single, composite transform to correct for head-motion and susceptibility distortions. These resampled BOLD time-series will be referred to as *preprocessed BOLD in original space*, or just *preprocessed BOLD*. The BOLD time-series were resampled into standard space, generating a *preprocessed BOLD run in ['MNI152NLin2009cAsym'] space*. First, a reference volume and its skull-stripped version were generated using a custom methodology of *fMRIPrep*. Several confounding time-series were calculated based on the *preprocessed BOLD*: framewise displacement (FD), DVARS and three region-wise global signals. FD and DVARS are calculated for each functional run, both using their implementations in *Nipype* (following the definitions by Power et al. 2014). The head-motion estimates calculated in the correction step were also placed within the corresponding confounds file. The confound time series derived from head motion estimates and global signals were expanded with the inclusion of temporal derivatives and quadratic terms for each (Satterthwaite et al. 2013). All resamplings can be performed with *a single interpolation step* by composing all the pertinent transformations (i.e. head-motion transform matrices, susceptibility distortion correction when available, and co-registrations to anatomical and output spaces). Gridded (volumetric) resamplings were performed using antsApplyTransforms (ANTs), configured with Lanczos interpolation to minimize the smoothing effects of other kernels (Lanczos 1964). Non-gridded (surface) resamplings were performed using mri_vol2surf (FreeSurfer).

Many internal operations of *fMRIPrep* use *Nilearn* 0.5.2 (Abraham et al. 2014, RRID:SCR_001362), mostly within the functional processing workflow. For more details of the pipeline, see the section corresponding to workflows in *fMRIPrep*'s documentation at (https://fmriprep.org/en/latest/workflows.html).

References for fMRI Preprocessing

Abraham, Alexandre, Fabian Pedregosa, Michael Eickenberg, Philippe Gervais, Andreas Mueller, Jean Kossaifi, Alexandre Gramfort, Bertrand Thirion, and Gael Varoquaux. 2014. "Machine Learning for Neuroimaging with Scikit-Learn." *Frontiers in Neuroinformatics* 8. https://doi.org/10.3389/fninf.2014.00014.

Avants, B.B., C.L. Epstein, M. Grossman, and J.C. Gee. 2008. "Symmetric Diffeomorphic Image Registration with Cross-Correlation: Evaluating Automated Labeling of Elderly and Neurodegenerative Brain." *Medical Image Analysis* 12 (1): 26–41. https://doi.org/10.1016/j.media.2007.06.004.

Behzadi, Yashar, Khaled Restom, Joy Liau, and Thomas T. Liu. 2007. "A Component Based Noise Correction Method (CompCor) for BOLD and Perfusion Based fMRI." *NeuroImage* 37 (1): 90–101. https://doi.org/10.1016/j.neuroimage.2007.04.042.

Cox, Robert W., and James S. Hyde. 1997. "Software Tools for Analysis and Visualization of fMRI Data." *NMR in Biomedicine* 10 (4-5): 171–78. https://doi.org/10.1002/(SICI)1099-1492(199706/08)10:4/5<171::AID-NBM453>3.0.CO;2-L.

Dale, Anders M., Bruce Fischl, and Martin I. Sereno. 1999. "Cortical Surface-Based Analysis: I. Segmentation and Surface Reconstruction." *NeuroImage* 9 (2): 179–94. https://doi.org/10.1006/nimg.1998.0395.

Esteban, Oscar, Ross Blair, Christopher J. Markiewicz, Shoshana L. Berleant, Craig Moodie, Feilong Ma, Ayse Ilkay Isik, et al. 2018. "FMRIPrep." *Software*. Zenodo. https://doi.org/10.5281/zenodo.852659.

Esteban, Oscar, Christopher Markiewicz, Ross W Blair, Craig Moodie, Ayse Ilkay Isik, Asier Erramuzpe Aliaga, James Kent, et al. 2018. "fMRIPrep: A Robust Preprocessing Pipeline for Functional MRI." *Nature Methods*. https://doi.org/10.1038/s41592-018-0235-4.

Fonov, VS, AC Evans, RC McKinstry, CR Almli, and DL Collins. 2009. "Unbiased Nonlinear Average Age-Appropriate Brain Templates from Birth to Adulthood." *NeuroImage* 47, Supplement 1: S102. https://doi.org/10.1016/S1053-8119(09)70884-5.

Gorgolewski, K., C. D. Burns, C. Madison, D. Clark, Y. O. Halchenko, M. L. Waskom, and S. Ghosh. 2011. "Nipype: A Flexible, Lightweight and Extensible Neuroimaging Data Processing Framework in Python." *Frontiers in Neuroinformatics* 5: 13. https://doi.org/10.3389/fninf.2011.00013.

Gorgolewski, Krzysztof J., Oscar Esteban, Christopher J. Markiewicz, Erik Ziegler, David Gage Ellis, Michael Philipp Notter, Dorota Jarecka, et al. 2018. "Nipype." *Software*. Zenodo. https://doi.org/10.5281/zenodo.596855.

Greve, Douglas N, and Bruce Fischl. 2009. "Accurate and Robust Brain Image Alignment Using Boundary-Based Registration." *NeuroImage* 48 (1): 63–72. https://doi.org/10.1016/j.neuroimage.2009.06.060.

Jenkinson, Mark, Peter Bannister, Michael Brady, and Stephen Smith. 2002. "Improved Optimization for the Robust and Accurate Linear Registration and Motion Correction of Brain Images." *NeuroImage* 17 (2): 825–41. https://doi.org/10.1006/nimg.2002.1132.

Klein, Arno, Satrajit S. Ghosh, Forrest S. Bao, Joachim Giard, Yrjö Häme, Eliezer Stavsky, Noah Lee, et al. 2017. "Mindboggling Morphometry of Human Brains." *PLOS Computational Biology* 13 (2): e1005350. https://doi.org/10.1371/journal.pcbi.1005350.

Lanczos, C. 1964. "Evaluation of Noisy Data." *Journal of the Society for Industrial and Applied Mathematics Series B Numerical Analysis* 1 (1): 76–85. https://doi.org/10.1137/0701007.

Power, Jonathan D., Anish Mitra, Timothy O. Laumann, Abraham Z. Snyder, Bradley L. Schlaggar, and Steven E. Petersen. 2014. "Methods to Detect, Characterize, and Remove Motion Artifact in Resting State fMRI." *NeuroImage* 84 (Supplement C): 320–41. https://doi.org/10.1016/j.neuroimage.2013.08.048.

Satterthwaite, Theodore D., Mark A. Elliott, Raphael T. Gerraty, Kosha Ruparel, James Loughead, Monica E. Calkins, Simon B. Eickhoff, et al. 2013. "An improved framework for confound regression and filtering for control of motion artifact in the preprocessing of resting-state functional connectivity data." *NeuroImage* 64 (1): 240–56. https://doi.org/10.1016/j.neuroimage.2012.08.052.

Tustison, N. J., B. B. Avants, P. A. Cook, Y. Zheng, A. Egan, P. A. Yushkevich, and J. C. Gee. 2010. "N4ITK: Improved N3 Bias Correction." *IEEE Transactions on Medical Imaging* 29 (6): 1310–20. https://doi.org/10.1109/TMI.2010.2046908.

Zhang, Y., M. Brady, and S. Smith. 2001. "Segmentation of Brain MR Images Through a Hidden Markov Random Field Model and the Expectation-Maximization Algorithm." *IEEE Transactions on Medical Imaging* 20 (1): 45–57. https://doi.org/10.1109/42.906424.
